# A novel functional gene delivery platform based on a commensal human anellovirus demonstrates transduction in multiple tissue types

**DOI:** 10.1101/2024.03.27.586964

**Authors:** Cato Prince, George Bounoutas, Bolu Zhou, Waseem Raja, Isabella Gold, Rianna Pozsgai, Parmi Thakker, Nicole Boisvert, Christopher Reardon, Stephanie Thurmond, Erin Ozturk, Rajendra Boggavarapu, Simeon Springer, Lovepreet Chahal, Maciej Nogalski, Tuyen Ong, Dhananjay Nawandar, Christopher Wright, Ashley Mackey, Geoffrey Parsons, Joseph Cabral

**Author notes:** Corresponding Author –.

## Abstract

*Anelloviridae* is a family of non-enveloped viruses with negative-sense, circular, single-stranded deoxyribonucleic acid (ssDNA) genomes that infect vertebrates and are a ubiquitous component of the human virome. Human anelloviruses evade induction of humoral immune responses and appear to be non-pathogenic. These properties, in conjunction with their enormous genomic diversity and wide tissue distribution, make anelloviruses compelling candidates as vectors for next-generation genetic medicines. Here we report the first gene delivery vector system based on a human commensal virus. This Anellovector is based on a virus of the *Betatorquevirus* genus. Production is enabled by the development of the Self-Amplifying Trans-complementation of a Universal Recombinant aNellovector (SATURN) system, which relies on a self-replicating plasmid to provide viral proteins in trans that drive replication and capsid-dependent packaging of vector genomes. The SATURN system also utilizes a Cre-lox-based recombination mechanism to generate single unit-sized circular genomes inside the MOLT-4 production cell line. We demonstrate that the SATURN system can package a vector genome from a single betatorquevirus with capsids from multiple betatorquevirus species, supporting the feasibility of establishing a novel vector platform that takes advantage of the remarkable diversity of anelloviruses. The Anellovector demonstrated function *in vitro* in retinal pigment epithelial (RPE) cells. The Anellovector also demonstrated durable *in vivo* function in the mouse eye for 9 months after subretinal administration, and achieved comparable gene expression to dose-matched adeno-associated virus 9 (AAV9) when transduced by the intracerebroventricular (ICV) route of administration. To our knowledge, this is the first report of a functional anellovirus-based gene therapy vector. Anellovectors have great potential to deliver safe, redosable, and potent therapeutics, helping to expand the reach of programmable medicines.

## INTRODUCTION

Viruses have evolved highly effective mechanisms for the delivery of genetic material into the nuclei of host cells, and it is this core function of viruses that has made them effective as vectors for gene therapy applications [Umair 2022]. Although viruses such as herpes simplex virus 1 (Vyjuvek for the treatment of epidermolysis bullosa) [Epstein 2023] and retroviruses (chimeric antigen receptor [CAR]-T cell therapies for cancer) [Charitidis 2023] have been used and even approved for various gene therapy applications, adeno-associated virus (AAV) has been the primary vector employed for the treatment of inherited monogenic disorders [Zhao 2021], which are individually rare but collectively affect >30 million Americans [Mendell 2021]. Several AAV-based gene therapies have been approved by the U.S. Food and Drug Administration (FDA) [Kohn 2023]. AAV vectors are engineered by removing the viral protein-coding sequences and substituting the therapeutic gene of interest as a payload [Wang 2019]. Approximately a dozen AAV serotypes are known to exist, with each exhibiting a unique tropism profile due to differing affinities for cell-surface attachment factors, receptors, or coreceptors [Li 2023]. While most AAV-based gene therapy trials have targeted the eye, liver, muscle, and CNS, other major organs such as the heart, kidney, and lung have been largely inaccessible to AAV-based gene therapies [Kuzmin 2021]. Limitations such as pre-existing neutralizing immunity and constraints on repeated dosing limit the broader clinical impact of AAV-based therapies [Mingozzi 2013, Mendell 2022]. At least 50% of humans have antibodies to AAV in their blood which can neutralize AAV-based gene therapies prior to transduction of target; these patients with neutralizing antibodies to AAV are often excluded from clinical trials [Wang 2019, Mendell 2022]. Following a single dose of an AAV-based gene therapy, a robust humoral immune response generally prevents re-administration. These limitations highlight the importance of expanding the gene therapy toolkit to include additional viral vectors with their own distinct properties.

*Anelloviridae* is a family of non-enveloped negative-sense, circular, ssDNA viruses known to infect vertebrate hosts [Biagini 2009, Butkovic 2023]. The first anellovirus identified was torque teno virus (TTV) in 1997 [Nishizawa 1997]. Within the anellovirus genera that have been identified across a broad range of vertebrates, only chicken anemia virus (CAV), of the *Gyrovirus* genus, is known to cause disease in its host [Fatoba 2019]. The remaining genera appear non-pathogenic and have seemingly evolved to coexist with their host species as commensal viruses [Virgin 2009, Kaczorowska 2020]. Anelloviruses comprise the majority of the known human virome [Ninomiya 2008, De Vlaminck 2013], and are grouped into three genera based on genome size: *Alpha*-, *Beta*-, and *Gammatorqueviruses*, also known as Torque teno virus (3.9 kb), Torque teno mini virus (2.8-2.9 kb), and Torque teno midi virus (3.2 kb), respectively [Biagini 2009, Varsani 2021, Arze 2021]. From infancy through adulthood, anelloviruses are detected in the blood and in virtually every tissue and organ of nearly 100% of people worldwide, remarkably without causing apparent disease [Freer 2018].

In addition to being the most prevalent virus in humans, anelloviruses are among the most genetically diverse. A vast number of unique anellovirus capsid sequences have been identified [Arze 2021], suggesting anelloviruses could be leveraged as a gene therapy vector platform with broad tropism for many sites of disease including those beyond the reach of current technologies. In addition, the well-documented persistence and replication of anelloviruses in humans without induction of humoral immune responses [Kaczorowska 2020] may minimize or avoid the problem of immunogenicity. In this regard, it has been suggested that anelloviruses evade host B-cell-mediated immunity by evolving at an antibody recognition site [Liou 2022]. Collectively, these properties of anelloviruses warranted the evaluation of an anellovirus-based gene therapy platform.

Here we report for the first time a completely novel gene therapy class termed Anellovectors. The Anellovector described here is based on a recently defined member of the *Betatorquevirus* genus, nrVL4619, that was recovered from human ocular tissue [Nawandar 2022]. Vectorization of anelloviruses, including nrVL4619, was made possible through development of the Self-Amplifying Trans-complementation of a Universal Recombinant aNellovector (SATURN) production platform, which relies on a self-replicating plasmid to provide viral proteins *in trans* that drive replication and packaging of vector genomes. Like most mature viral vector systems, the vector genome itself lacks viral coding sequences and only carries therapeutic or reporter transgenes and the viral cis elements required for DNA replication and packaging. Using the SATURN platform, we can produce capsid protein-dependent particles that encapsidate circular ssDNA vector genomes. We characterized these particles by DNase-protected quantitative polymerase chain reaction (qPCR), next-generation sequencing, and electron microscopy (EM). Furthermore, we demonstrate packaging of a vector genome from a single anellovirus with capsids from multiple anellovirus species, suggesting a universal vector platform that takes advantage of the diversity of anelloviruses can be built*. In vitro* transduction was validated by detection of an enhanced green fluorescent protein (eGFP) reporter and detection of vector genomes in nuclei by *in situ* hybridization. *In vivo* expression of an eGFP reporter was observed in both ocular and central nervous system (CNS) mouse models, demonstrating the potential for in-life use of Anellovectors in delivering therapeutic payloads.

## RESULTS

### Development of the SATURN platform for the production of Anellovectors

Anellovirus genomes consist of circular ssDNA. Shortly after entry to the nucleus, it is presumed that the viral genome forms double-stranded DNA (dsDNA) replication intermediates to facilitate viral gene expression and act as templates for the replication of progeny viral genomes. Anelloviruses, unlike AAVs or lentiviruses, do not have inverted terminal repeats (ITRs) that define genomic boundaries and enable replication intermediates from a plasmid context; therefore, an input Anellovector genome must be able to form a circular dsDNA replication intermediate of the genomic unit size during production. We constructed a vector plasmid that utilizes the Cre-lox recombination system [Hoess 1990] to produce unit-sized replication intermediate vectors from a plasmid backbone inside the production cell **(Figure 1A).** This is accomplished by flanking a cassette, comprised of the entire nrVL4619 betatorquevirus noncoding region (NCR) followed by the full cytomegalovirus (CMV) promoter driving an eGFP transgene, with an upstream lox71 and downstream lox66. In this configuration, recombination by the Cre recombinase will yield a single genomic unit-size (2839 bp) dsDNA molecule with a loxP site and a 2613-bp plasmid backbone that contains a lox72 site. A woodchuck hepatitis virus posttranscriptional regulatory element (WPRE) was added following the eGFP cassette to enhance expression of the transgene. We refer to this vector as ANV.eGFP.

**Figure 1:**
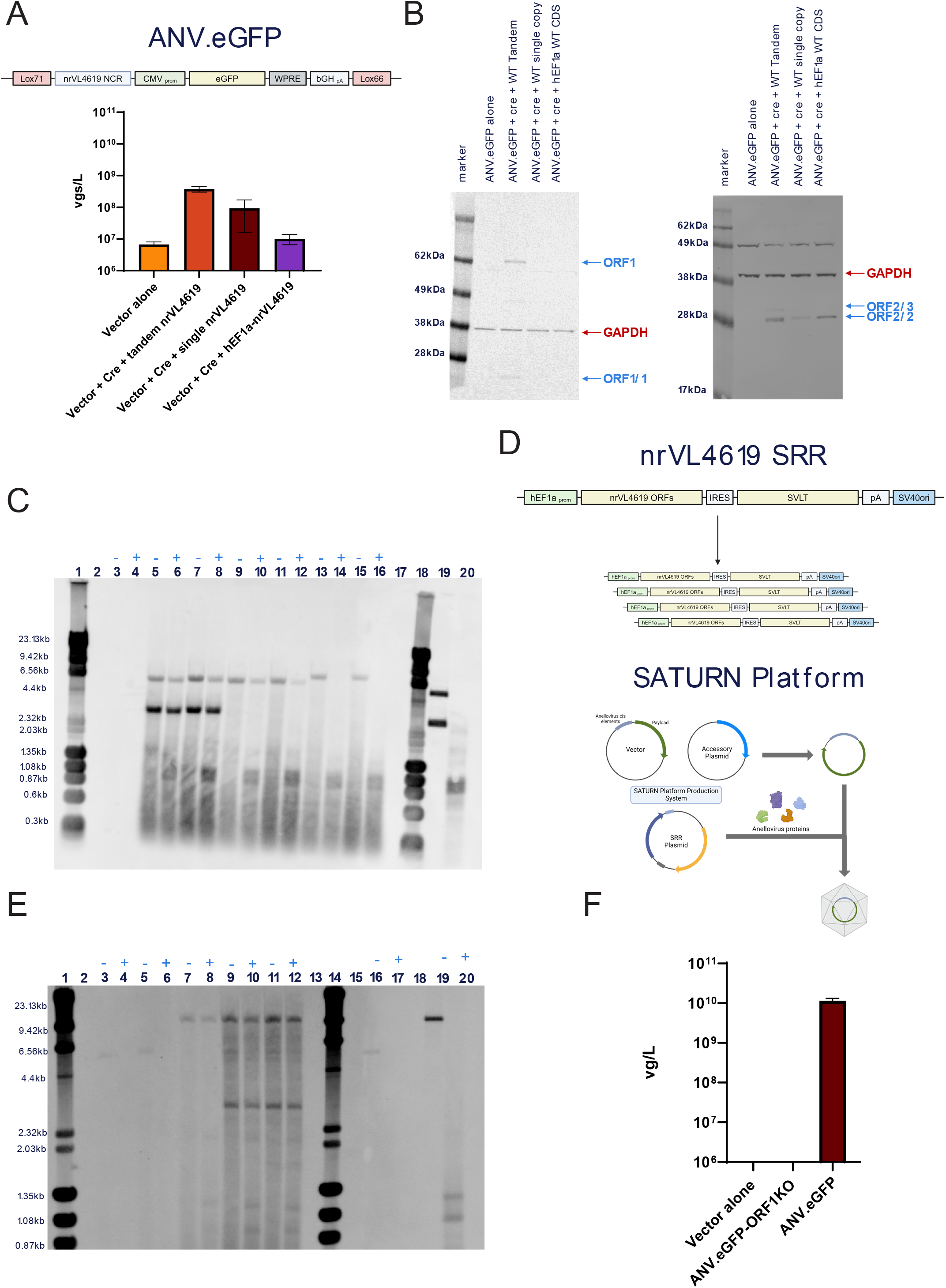
Development of the SATURN system. (A) (Top) Schematic representation of the Anellovector comprising the nrVL4619 NCR, CMV promoter, eGFP, WPRE, and bGH polyA flanked by the mutant lox sites, Lox71 and Lox66. (Bottom) Comparison of DNase-protected vector DNA (using eGFP (4) primer-probe set) when viral proteins are supplied by plasmids of nrVL4619 WT single or tandem or hEF1α-nrVL4619-CDS; data reported as vector genomes per liter of production culture (vgs/L) and presented as mean (n=3) ± SEM. (B) Immunoblots probing for GAPDH and the viral proteins (ORF1 and ORF1/1 left; ORF2, ORF2/2, and ORF2/3 right) from nrVL4619 WT and hEF1α-driven contexts in MOLT-4 cells. The blue arrows denote the expected sizes of the viral proteins. Red arrow denotes the GAPDH loading control. (C) Southern blot probing against nrVL4619 CDS sequences for conditions outlined in Table 1 (Top). The plus and minus signs indicate whether the sample was digested with DpnI to remove input plasmid DNA. See Table 1 (top) for lane identities and restriction enzyme digests. Sizes of expected digestion products: nrVL4619 tandem 5479bp + 2876bp (single unit genome); nrVL4619 single copy 5479; hEF1a-nrVL4619-CDS 6636. D) Schematic diagram of the nrVL4619 SRR (Top) and the SATURN system (bottom). The SRR contains a hEF1α promoter driving the co-expression of the nrVL4619 and SV40 large T antigen. The downward arrow and multiple smaller SRRs denote the self-amplification of the plasmid by large T antigen and the SV40 origin of replication. The SATURN system comprises a three-plasmid transfection which circularizes the vector genome out of a plasmid (“Vector”) through the expression of Cre (“Accessory plasmid”) and the SRR plasmid supplying anellovirus proteins for replication and packaging. (E) Southern blot probing against nrVL4619 CDS and eGFP transgene sequences (multiplex) for conditions outlined in Table 1 (bottom). The plus and minus signs indicate whether the sample was digested with DpnI to remove input plasmid DNA. See Table 1 (bottom) for lane identities and restriction enzyme digests. Sizes of expected digestion products: ANV.eGFP plasmid 5452bp; ANV.eGFP single unit replication intermediate 2839bp; nrVL4619 SRR single digestion 9391bp; ORF1-KO SRR single digestion 9391bp. (F) Quantitation of ANV.eGFP DNase-protected vector DNA recued with either the nrVL4619 SRR or ORF1-KO SRR using eGFP (4) primer-probes. Data presented as mean (n=3) ± SEM.

**Table 1.**

Fig 1C key - Probe: nrVL4619 genomic
| Lane | Condition | Enzyme(s) |
| --- | --- | --- |
| 1 | Mixed Ladder: $\lambda$ DNA- HindIII Digest + ϕX174 DNA- HaeIII Digest | |
| 2 | empty |  |
| 3 | ANV.eGFP alone | AgeI- HF |
| 4 | ANV.eGFP alone | AgeI- HF + DpnI |
| 5 | ANV.eGFP + Cre + nrVL4619 Tandem | EcoRV- HF |
| 6 | ANV.eGFP + Cre + nrVL4619 Tandem | EcoRV- HF + DpnI |
| 7 | nrVL4619 Tandem alone | EcoRV- HF |
| 8 | nrVL4619 Tandem alone | EcoRV- HF + DpnI |
| 9 | ANV.eGFP + Cre + nrVL4619 single copy | EcoRV- HF |
| 10 | ANV.eGFP + Cre + nrVL4619 single copy | EcoRV- HF + DpnI |
| 11 | nrVL4619 single copy alone | EcoRV- HF |
| 12 | nrVL4619 single copy alone | EcoRV- HF + DpnI |
| 13 | ANV.eGFP + Cre + hEF1a-nrVL4619 CDS | EcoRV- HF |
| 14 | ANV.eGFP + Cre + hEF1a-nrVL4619 CDS | EcoRV- HF + DpnI |
| 15 | hEF1a-nrVL4619 CDS alone | EcoRV- HF |
| 16 | hEF1a-nrVL4619 CDS alone | EcoRV- HF + DpnI |
| 17 | empty |  |
| 18 | Mixed Ladder: $\lambda$ DNA- HindIII Digest + ϕX174 DNA- HaeIII Digest | |
| 19 | 1e+09 copies of nrVL4619 single copy plasmid (control plasmid digest) | EcoRV- HF |
| 20 | 1e+09 copies of nrVL4619 single copy plasmid (control plasmid digest) | EcoRV- HF + DpnI |

Fig 1E key - Probe: nrVL4619 genomic + eGFP
| Lane | Condition | Enzyme(s) |
| --- | --- | --- |
| 1 | Mixed Ladder: $\lambda$ DNA- HindIII Digest + ϕX174 DNA- HaeIII Digest | |
| 2 | empty |  |
| 3 | ANV.eGFP alone | AgeI- HF |
| 4 | ANV.eGFP alone | AgeI- HF + DpnI |
| 5 | ANV.eGFP + Cre | AgeI- HF |
| 6 | ANV.eGFP + Cre | AgeI- HF + DpnI |
| 7 | nrVL4619 SRR alone | AgeI- HF |
| 8 | nrVL4619 SRR alone | AgeI- HF + DpnI |
| 9 | ANV.eGFP + Cre + nrVL4619 SRR | AgeI- HF |
| 10 | ANV.eGFP + Cre + nrVL4619 SRR | AgeI- HF + DpnI |
| 11 | ANV.eGFP + Cre + ORF1-KO SRR | AgeI- HF |
| 12 | ANV.eGFP + Cre + ORF1-KO SRR | AgeI- HF + DpnI |
| 13 | empty |  |
| 14 | Mixed Ladder: $\lambda$ DNA- HindIII Digest + ϕX174 DNA- HaeIII Digest | |
| 15 | empty |  |
| 16 | 1e+09 copies of ANV.eGFP | AgeI- HF |
| 17 | 1e+09 copies of ANV.eGFP | AgeI- HF + DpnI |
| 18 | empty |  |
| 19 | 1e+09 copies of nrVL4619 SRR | AgeI- HF |
| 20 | 1e+09 copies of nrVL4619 SRR | AgeI- HF + DpnI |

We recently reported the development of a wild-type (WT) virus production system based on the use of MOLT-4 cells [Minowada 1972], a T lymphoblast cell line [Nawandar 2022]. We now utilize MOLT-4 cells as a model system for producing Anellovector. To supply viral proteins *in trans*, we tested multiple strategies, including co-transfection with WT genomes in both single and tandem configurations as well as designing an expression construct in which the nrVL4619 coding sequences were driven by a human eukaryotic translation elongation factor 1 α (hEF1α) promoter. While both the single and tandem WT genome constructs of nrVL4619 were able to produce detectable vector as measured by DNase-protected qPCR from linear iodixanol gradient-purified material, the hEF1α-nrVL4619 construct did not produce vector above background **(Figure 1A).** However, both WT genome constructs produced detectable levels of WT DNase-protected genomes **(Supplementary Figure 1A),** suggesting they may produce WT virion contaminants. Interestingly, all three viral protein expression plasmids produced detectable quantities of the ORF2/2 accessory protein, but only the nrVL4619 tandem plasmid could produce detectable levels of the ORF1 capsid protein (Figure 1B). The lack of a band for ORF1 in the nrVL4619 single genome copy and hEF1α-nrVL4619 samples may be due to weak sensitivity of the ORF1 antibody. A band in the ORF2 antibody blot was detected at 49 kDa in all lanes and is likely the result of cross-reactivity with a host cell protein. Despite the hEF1α-nrVL4619 construct not driving encapsidation of the ANV.eGFP vector DNA, it could drive replication of a single genomic unit-sized (2839 bp) vector dsDNA replication intermediate inside the cell, as detected by a DpnI restriction-enzyme protected Southern assay **(Supplementary Figure 1B and Supplementary Table 1).** DpnI recognizes the methylated adenine of GATC sequences that have been replicated in *E. coli* and thus digests input DNA, while DNA that has been replicated in eukaryotic cells will be DpnI-resistant [Rao 1988, Wilson 2012]. Notably, while the nrVL4619 single and tandem genome plasmids were capable of self-replication, as seen in both the 2876-bp single unit genome and 5479-bp plasmid bands, the hEF1α-nrVL4619 plasmid was not **(Figure 1C and Table 1).** Taken together, these observations suggest that Cre recombinase forms a single genomic unit-sized vector replication intermediate inside the MOLT-4 cells, and gene products from the single and tandem nrVL4619 genome constructs can drive ssDNA conversion of replication intermediates and encapsidation of ANV.eGFP vector genomes. However, they produce WT virion contaminants. Notably, the plasmids that produced DNase-protected copies of ANV.eGFP were themselves capable of self-replication.

When probing for vector and WT genomes by Southern blot, we observed a potential correlation between the self-replication of the plasmids encoding trans viral elements and the production of packaged vector as measured by DNase-protected genomes **(Figures 1A and 1C).** These observations suggested that the WT viral genome constructs could drive production of vector in the MOLT-4 system, but the presence of WT contaminants necessitated development of another source for trans complementation with viral proteins, potentially with the capacity for self-replication. Thus, in order to develop a WT-free vector production system for Anellovectors, we investigated the possibility of using a rescue plasmid with a self-replicating architecture in hopes of enhancing the expression of anellovirus proteins in trans while preventing WT contamination of the vector product. To this end, we developed a self-replicating rescue plasmid (SRR) that exploits the simian virus 40 (SV40) replication machinery [Collins 1992] to drive self-amplification of the rescue plasmid template. The SRR consists of the nrVL4619 coding sequences in their native configuration driven by a hEF1α promoter followed by the internal ribosome entry site 2 (IRES2) of the encephalomyocarditis virus (EMCV), which enables translation of a SV40 large T antigen cassette **(Figure 1D).** Expression of the large T antigen initiates replication of the plasmid from a downstream SV40 origin (Ori) of replication. By co-expressing large T antigen with the viral proteins, a forward feedback loop is established whereby more plasmid copies are replicated by the large T antigen, which in turn expresses more viral proteins and large T antigen. The SRR was combined in a triple transfection with the lox-site flanked Anellovector and a Cre expression plasmid **(Figure 1D).** The resulting production system was termed the SATURN platform.

To test the nrVL4619 SRR construct, we first confirmed expression of viral proteins as detected by immunoblot **(Supplementary Figure 1C).** Next, we performed a triple transfection with the full SATURN platform into MOLT-4 cells which resulted in recovery of DNase-protected vector genomes after linear iodixanol gradient purification **(Figure 1E).** The DNase-protected signal for WT DNA dropped below the limit of quantitation **(Supplementary Figure 1D).** To verify that the DNase-protected vector genome signal was dependent on capsid protein, we replaced the codon for phenylalanine 46 (F46) of ORF1 with a premature stop codon in the nrVL4619 SRR (ORF1-KO SRR), specifically disrupting translation in a sequence unique to the ORF1 open reading frame. The ORF1-KO SRR produced accessory protein ORF2/2, but it did not produce detectable levels of ORF1 by immunoblot **(Supplementary Figure 1C).** While the ORF1-KO SRR was capable of driving replication of a 2839-bp single genomic unit-sized vector replication intermediate **(Figure 1E),** it failed to produce DNase-protected signal for ANV.eGFP **(Figure 1F).** Both of the SRR variants could drive self-replication of the plasmid as demonstrated by the presence of a 9391-bp DpnI-resistant digestion product **(Figure 1E).** These data support the assertion that the SATURN platform’s self-replicating architecture can provide the viral components required to produce a WT-free ANV.eGFP vector in a capsid-dependent manner using a MOLT-4 cell system.

### Anellovectors package circular ssDNA into 30-nm vector particles

The SATURN platform can produce capsid-dependent DNase-protected qPCR signal for ANV.eGFP vector **(Figure 1);** however, these data alone are insufficient to validate the production of fully encapsidated vector. To further characterize the protected vector DNA, we performed a DNase-protected qPCR assay on linear iodixanol gradient-recovered material that used 4 primer-probe sets that span the eGFP cassette **(Figure 2A).** While the vector alone negative control and the ORF1-KO SRR conditions showed only background signal, the ANV.eGFP with SRR conditions showed a similar DNase-protected signal for all 4 primer-probe sets suggesting the full transgene was intact. We next investigated whether the detected DNA was circular in nature. When Cre recombines the lox71 and lox66 variant sites, it results in a loxP and lox72 site on the two daughter DNA molecules [Albert 1995]. We validated qPCR primer-probe sets specific for lox72 and loxP to confirm that the DNase-protected ANV.eGFP DNA produced by the SATURN platform is specific for the loxP site **(Supplementary Figure 2A and 2B).** The signal for the vector-specific loxP site and the eGFP signal were nearly identical, demonstrating that the ANV.eGFP protected by the ORF1 protein was circular and intact **(Figure 2B).**

**Figure 2:**
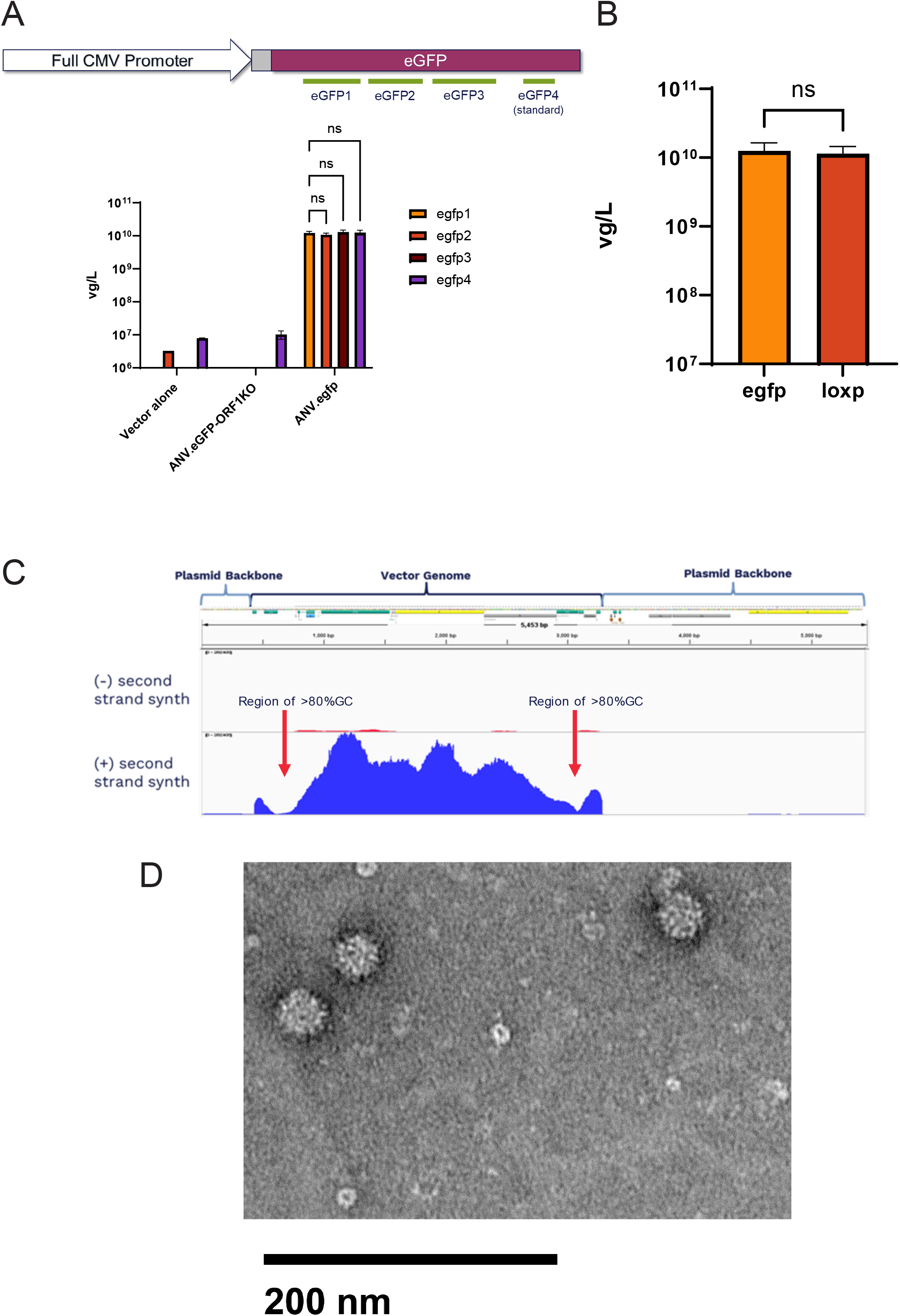
Anellovectors package ssDNA genomes into 30-nm particles. (A) (Top) Schematic diagram of the eGFP qCPR tiling assay; green bars below egfp represent four different regions of egfp probed for titer. (Bottom) Comparison via qPCR of DNase-protect ANV.eGFP DNA between four egfp primer-probe sets. Data presented as mean (n=3) ± SEM; ns (two-way ANOVA). (B) Comparison of DNase-protected ANV.eGFP DNA using probes targeting either the egfp gene (egfp (4) primer-probe set) or the loxP site. Data presented as mean (n=3) ± SEM; ns: p > 0.05 (two-tailed t-test). (C) Coverage plot of sequenced vector genomes mapped to diagram of vector plasmid. Minus sign indicates coverage of samples not processed with an additional second strand synthesis step; plus sign indicates coverage of samples processed with a second strand synthesis step. Red arrows denote areas of high GC content. (D) Electron microscopy of vector particles at 30,000 × magnification; 200-nm bar for scale.

Next, we investigated whether the ORF1-protected DNA was single-stranded or double-stranded and if plasmid backbone was also encapsidated. We produced purified ANV.eGFP using the SATURN platform coupled with linear iodixanol gradient purification, buffer exchange, and concentration. The resulting vector preparation was analyzed by next-generation sequencing (NGS). Any reads mapped to the vector genome confirmed the DNA was single-stranded in nature, as sequencing is dependent on second strand synthesis during library preparation **(Figure 2C).** We observed that 97.5% of mapped reads aligned to the vector cassette flanked by lox sites. There was a reduction in reads through two areas of >80% GC content, with one located in the nrVL4619 NCR and the other in the polyadenylation signal for the eGFP cassette. Notably, fewer than 1% of reads mapped to the plasmid backbone, suggesting anellovirus encapsidation is specific for sequences that contain the nrVL4619 NCR. We next examined SATURN-produced vector by EM to determine if virus-like particles (VLPs) are produced. We observed the presence of symmetrical icosahedral particles approximately 30 nm in size **(Figure 2D).** These VLPs are nearly identical in appearance to WT nrVL4619 virions produced by a WT tandem construct in a manner previously described [Nawandar 2022] **(Supplementary Figure 2C).** Taken together, these data demonstrate the SATURN platform can produce WT-free ANV.eGFP vector particles that encapsidate circular ssDNA in a capsid-dependent manner.

### Multiple anellovirus lineages can package an nrVL4619-based vector

The SATURN platform was designed to take advantage of the diversity of anelloviruses by utilizing a self-replicating architecture in which the viral coding sequences can be replaced by that of any anellovirus to replicate and package a universal vector template DNA **(Figure 3A).** To test this design, we produced SRR plasmids containing the coding sequences of TTMV-LY1 (NC_020498.1) and TTMV-LY2 (JX134045.1) [GalmLs 2013], which exhibit 38.1% and 31.9% amino acid similarity to the ORF1 of nrVL4619, respectively. The LY1 and LY2 SRRs were each co-transfected with an ANV.eGFP vector construct and a Cre expression plasmid. Both the LY1 and LY2 SRRs were capable of cross-packaging the ANV.eGFP vector genome, albeit with varying degrees of productivity **(Figure 3B).** The LY2-based SRR could produce DNase-protected titers near the level of the nrVL4619 SRR, while the level of production by the LY1-based SRR was close to 10-fold lower. Proteins from the LY2-based SRR were able to drive replication of the nrVL4619-based ANV.eGFP vector DNA as determined by Southern assay **(Figure 3C);** however, the LY1-based SRR did not produce levels of replicated DNA that were detectable. This corresponds with the 10x lower yield of vector recovered from the iodixanol gradient, and this may be because the amount of DNA was beneath the lower limit of detection for our assay. Both of these observations support the conclusion that proteins from distantly related anelloviruses can be used to cross-replicate and package nrVL4619-based vector DNA and highlight the potential of this construct as a universal vector template for future Anellovectors.

**Figure 3:**
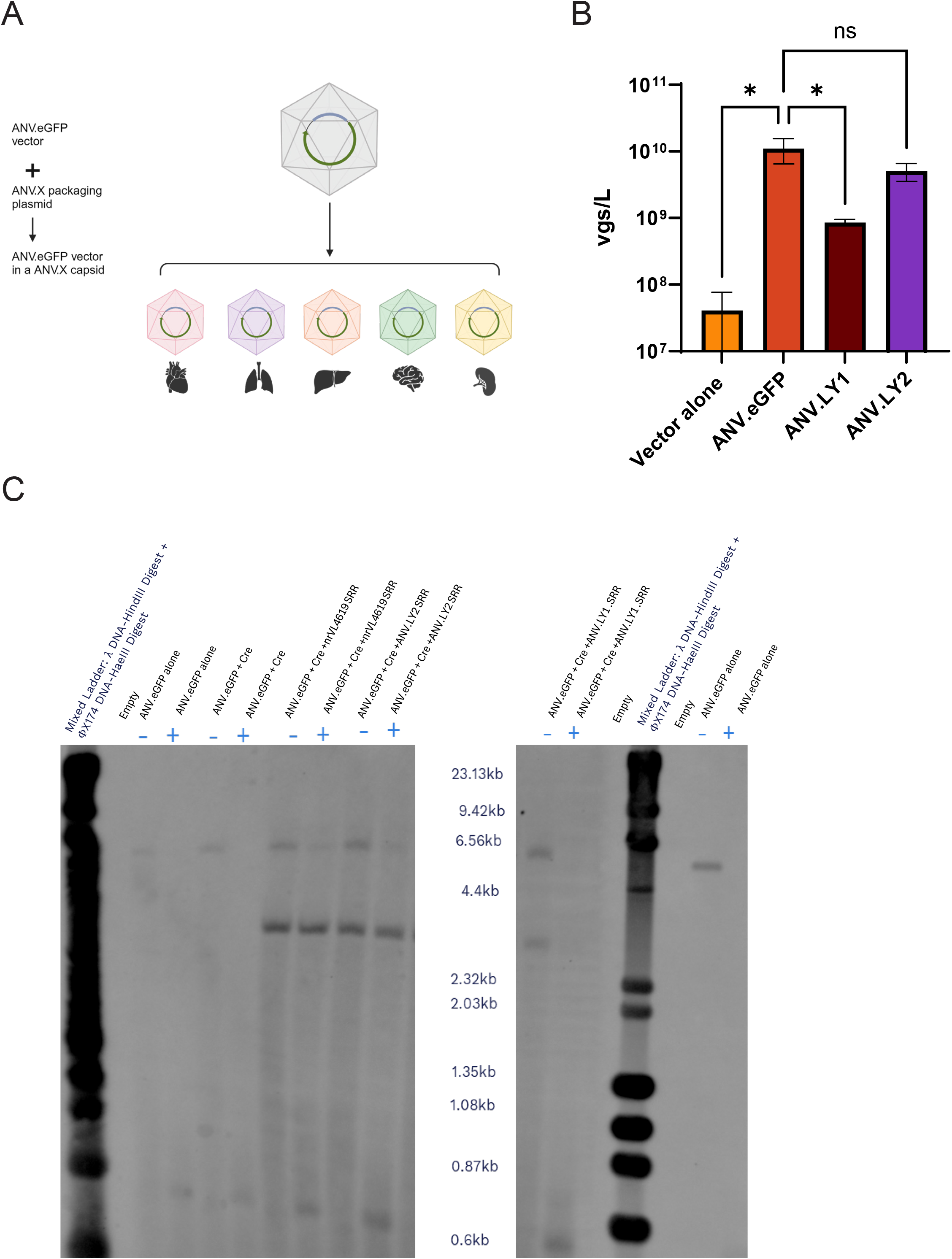
Cross-packaging of an nrVL4619-based vector into different anellovirus capsids. (A) Diagram representing the discovery of anellovirus species from various tissues of origin and their utilization to encapsidate the vector of ANV.eGFP. (B) Comparison of productivity as measured by DNase-protected qPCR detecting ANV.eGFP DNA packaged into its own capsid versus being packaged into ANV.LY1 or ANV.LY2. Data presented as mean (n=3) ± SEM; *: p< 0.05 (one-way ANOVA); ns: p > 0.05 (one-way ANOVA with Brown-Forsythe test). (C) Southern blot probing against eGFP transgene. All samples were digested with the AgeI restriction enzyme to linearize DNA. The plus and minus signs indicate whether the sample was digested with DpnI to remove input plasmid DNA. Expected band sizes: ANV.eGFP plasmid 5452bp; ANV.eGFP replication intermediate 2839bp.

### Anellovector demonstrates transduction of RPE cells in culture

The sequence of the nrVL4619 betatorquevirus was originally recovered in patient-derived ocular tissue, specifically the retinal pigment epithelium [Nawandar 2022]. To determine if the vectorized form of nrVL4619, ANV.eGFP, would retain tropism for cells derived from its tissue of origin, we used a culture model of induced pluripotent stem cell (iPSC)-derived retinal pigment epithelial cells (RPEs) which displayed the hallmark cobblestone-like morphology with pigmentation **(Figure 4A).** Following transduction with ANV.eGFP, we utilized *in situ* hybridization with a probe specific for the vector genome. We observed vector DNA in the nucleus within 24 hours that persisted for 7 days post-transduction **(Figure 4B).** Additionally, at 7 days post-transduction we detected eGFP transcripts by reverse transcription digital droplet PCR (RT-ddPCR) **(Figure 4C)** as well as by *in situ* hybridization, probing samples at 7 to 28 days post-transduction **(Figure 4D).** The presence of eGFP was confirmed by immunostaining cells for eGFP in conjunction with the *in situ* hybridization for eGFP mRNA detection, with eGFP-positive cells also positive for mRNA signal throughout the cytoplasm of the cell **(Figure 4D).** These experiments confirmed that the nrVL4619-based ANV.eGFP vector retained the tropism of its parental anellovirus for RPE.

**Figure 4:**
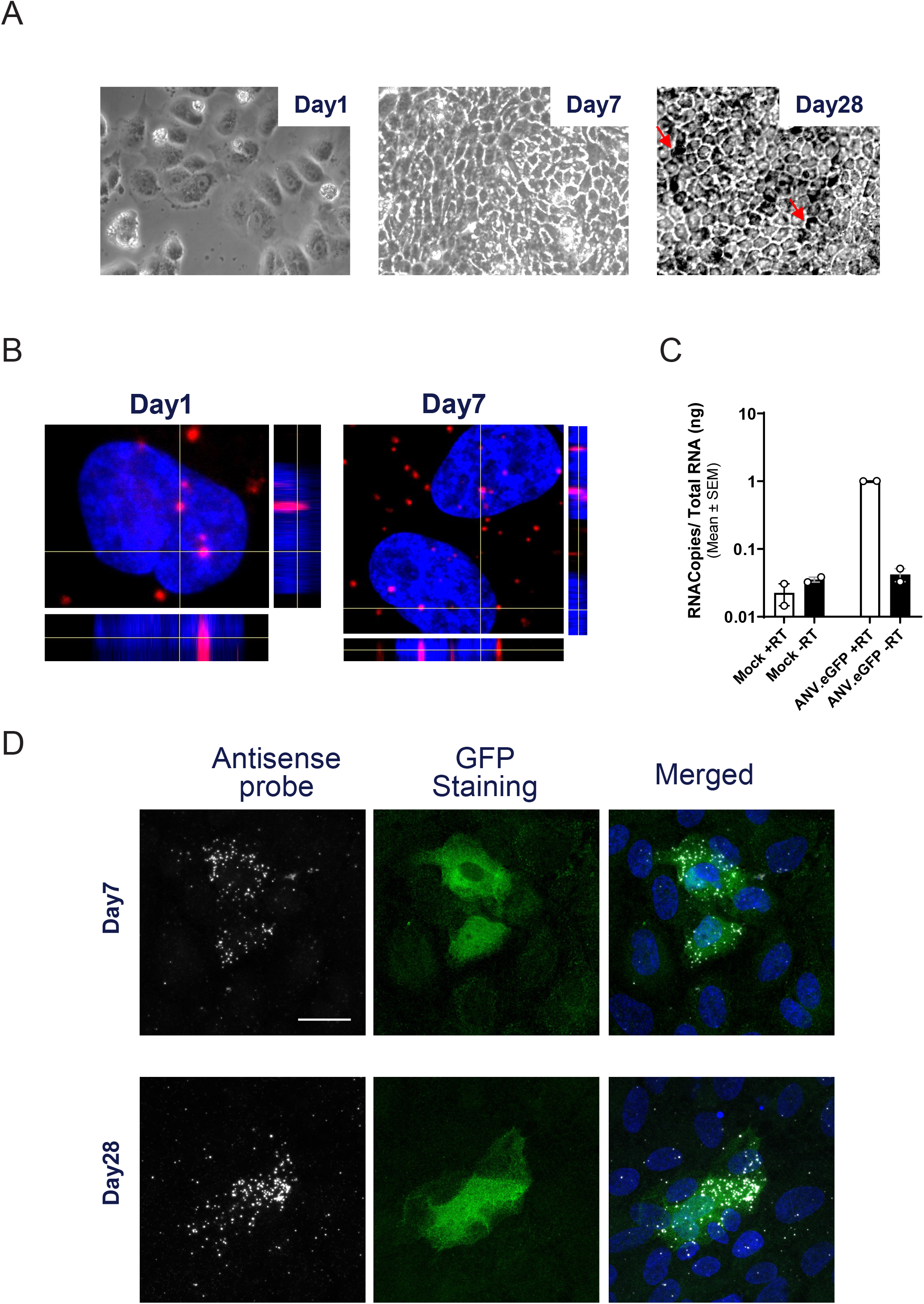
ANV.eGFP transduces RPE cells in culture. (A) Phase contrast images of Retinal pigment epithelial (RPE) on days 1, 7, and 28 after transduction at 40X magnification. On day 28 different levels of pigment structure were observed in the culture (red arrows). (B) Post RNase treatment in-situ hybridized cross-sectional view of RPE nuclei stained with DAPI (blue) and a probe to detect negative sense vector DNA followed by fluorophores TSA vivid 650 (red), the cross-section view showing the nuclear volume (MOI of 2K [day 1] and 3K [day 7] vg/cell) at a magnification of 63X and a scale bar represent 5 um. (C) RT-ddPCR plot of RPE on mock and ANV.eGFP treated samples with and without reverse transcriptase in biological duplicate on day 7 (MOI 2K vg/cell). (D) Immunostaining and in-situ hybridization of RPE on Day7 (MOI 6K) and Day 28 (MOI 0.9K). The antisense probe was used to detect the transcript level at the cellular resolution (left panels),the GFP antibody staining was used to detect the expression of protein (middle panels). The third panels show the merged images at a magnification of 63X. The scale bar represents 20 um.

### Subretinal delivery of an Anellovector demonstrated long-term durability and a favorable safety profile in a murine ocular model

ANV.eGFP demonstrated tropism for iPSC-derived RPE cells in culture; however, it still remained to investigate ANV.eGFP tropism and function in an ocular model *in vivo*. To evaluate the *in vivo* tropism, durability, and safety of the Anellovector in the eye, C57BL/6J mice were subjected to subretinal injections of ANV.eGFP at a dose of 1E8 vg **(Figure 5A).** Experimental eyes were collected at 3-, 6-, and 9-months post-injection, and the posterior eye cup (PEC), comprised of RPE cells, was isolated for tropism and durability analyses. The durability of ANV.eGFP was demonstrated by the detection of eGFP DNA and mRNA expression in the PEC over the 9-month period **(Figure 5B and C).** Although there was a 30x decrease in ANV.eGFP DNA copies observed in the PEC from 3 to 6 months, DNA levels remained relatively stable from 6 to 9 months, exhibiting only a 3x decrease. Despite the decline in DNA copies over time, the eGFP transgene expression remained stable from 3 to 9 months. The durable expression of the ANV.eGFP vector over 9 months demonstrates that the Anellovector can transduce its parental virus tissue of origin after subretinal injection and corroborates our *in vitro* observations.

**Figure 5.**
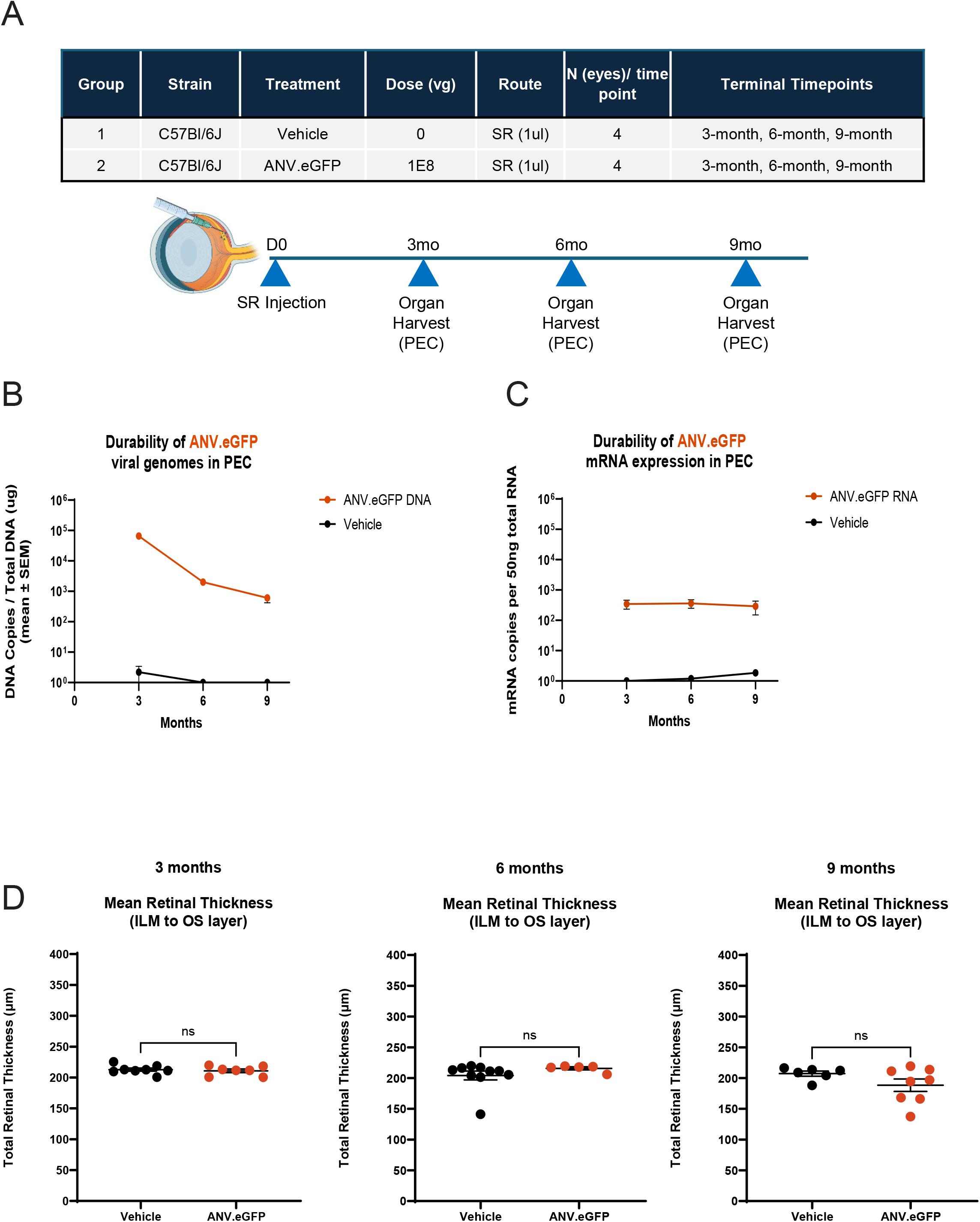
ANV.eGFP viral DNA and RNA durability in posterior eye cup. (A) Table describing strain of mouse, treatments, vector dose, route of administration, numbers of animals per group, and time points for the *in vivo* study. Bottom panel shows a schematic of a mouse eye’s anatomy and study design. (B) Average vector genome copies present in PEC, as assessed by qPCR in the harvested DNA from mouse eyes injected subretinally with either vehicle or 1E+8 vg of Anellovector. n = 4 eyes/group. (C) Absolute mRNA copies present in PEC, as assessed by RT-ddPCR in the eyes injected subretinally with either vehicle or 1E+8 vg of Anellovector. n = 4 eyes/group. (D) Mean retinal thickness measurements of OCT images from C57Bl/6J female mice at 3-, 6-and 9-months post-subretinal injection of either ANV.eGFP vector or vehicle. Measurements represent the distance between the ILM and the OS layers on each eye (n=5-10 eyes/timepoint) at specified intervals. Error bars indicate SEM.

To assess the safety of ANV administration *in vivo*, we examined the health and anatomical structure of each study eye using optical coherence tomography (OCT). OCT is a non-invasive, quantitative diagnostic tool commonly used in ophthalmology to evaluate eye health, particularly the retina, allowing for detailed examination of retinal structure [Stuarenghi 2014, Zeppieri 2023]. For our analysis, we focused on measuring retinal thickness, employing a customized approach, measuring from the internal limiting membrane (ILM) to the outer photoreceptor segment (OS). This method provided sufficient information to assess retinal health and structure sequentially at 3-, 6-, and 9-month intervals following subretinal injection of vehicle or ANV.eGFP vector. No retinal toxicity of ANV.eGFP was observed through 9 months, as indicated by the measurement of total retinal thickness in ANV.eGFP-injected eyes compared to vehicle-injected eyes **(Figure 5D).**

### Anellovector demonstrates similar transduction to AAV9 vector following ICV administration

While durable transgene expression was observed *in vivo* in the mouse eye, we were also interested in exploring the potential for Anellovector tropism outside the eye. For this purpose, we evaluated ANV transduction in the CNS of mice. ANV.eGFP or AAV9.eGFP (positive control) was administered by ICV injection through the left and right lateral ventricles in the brain (Figure 6A). ANV.eGFP showed comparable levels of eGFP vector DNA compared to AAV9.eGFP vector DNA in the brain at 21 days post-transduction, demonstrating comparable infectivity of ANV.eGFP versus AAV9.eGFP (Figure 6B). Additionally, consistent with these DNA levels, mRNA levels of ANV.eGFP and AAV9.eGFP in the brain were comparable, indicating similar transduction efficiency of ANV.eGFP compared to AAV9.eGFP (Figure 6C). To further examine the transduction of ANV.eGFP in the brain, sections from the brains of mice that received either ANV.eGFP or AAV9.eGFP were stained with an eGFP antibody (Figure 6D). A significant eGFP signal was observed in the hippocampus of brain sections from ANV.eGFP-treated mice, and this signal was comparable to the AAV9.eGFP-treated group. Interestingly, the cells positive for ANV.eGFP showed intact morphology with extended axons. In contrast, cells in the AAV9.eGFP group showed dispersed and disrupted axon morphology (Figure 6D). Taken together, these data demonstrate the Anellovector has tropism for cells in the CNS and suggests it can express its transgene to similar levels as an AAV9-based vector.

**Figure 6.**
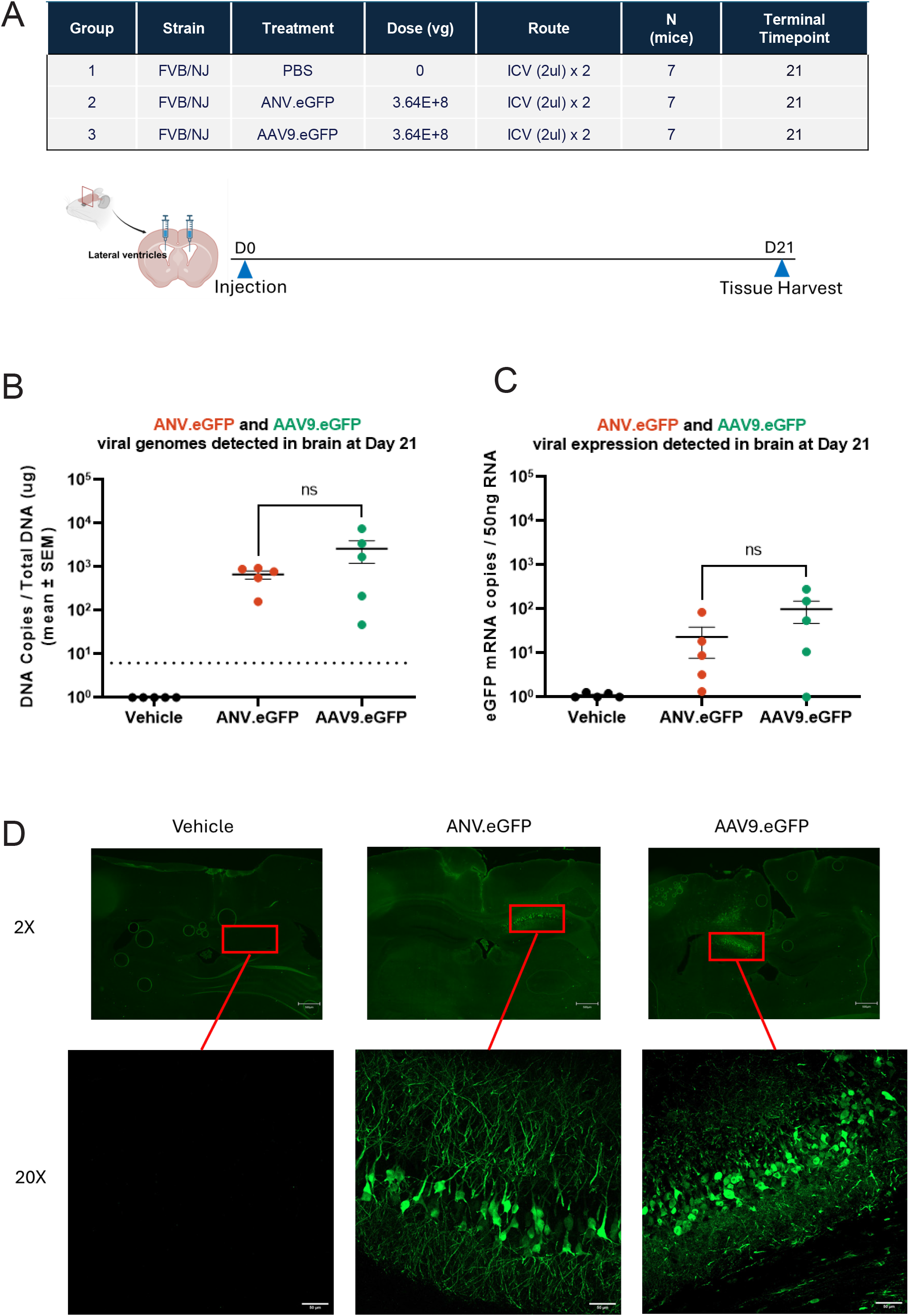
ANV.eGFP shows comparable transduction efficiency to AAV9.eGFP in the brain. (A) Table describing strain, treatment, vector doses, route of administration, animal numbers and terminal timepoint for *in vivo* study. The bottom schematic represents study design with ICV injection in mouse at Day 0 and brains harvested at Day 21. (B) eGFP DNA copies by qPCR in one hemisphere of the brains from mice that received either ANV.eGFP or AAV9.eGFP ICV injections (n=5/group). Statistical analyses were conducted using one-way ANOVA. (C) Absolute mRNA copies by RT-ddPCR in one hemisphere of the brains from mice that received either ANV.eGFP or AAV9.eGFP ICV injections (n=5/group). Statistical analyses were conducted using one-way ANOVA. (D) Representative immunofluorescence imaging for eGFP using eGFP antibody in the hippocampus of the brain sections from mice that received either ANV.eGFP or AAV9.eGFP. Magnification 2X, scale bar 500 µm; magnification 20X, scale bar 50 µm.

## DISCUSSION

There is a current unmet need for novel gene therapy vectors that build on the successes of existing gene therapy vectors while overcoming their limitations. Candidate technologies under investigation include viral and non-viral vectors. Non-viral vectors, e.g., lipid nanoparticles and polymeric-based vehicles, can be designed with attractive features such as high gene-loading capacity [Wang 2023]; however, technical questions regarding the *in vivo* stability of non-viral vector-based gene therapies and their potential for off-target effects remain unanswered [Rajawat 2019]. Clinical translation of these approaches has not been realized. All approved gene therapies and most gene therapies in clinical trials employ viral vectors [Arabi 2022], which represent a clinically validated strategy for introducing therapeutic genes into patients. What has been missing from the field of gene therapy is an alternative viral vector platform with wider applicability to heretofore intractable diseases and lower risks of immunogenicity and toxicity [Shen 2022, Srivastava 2023]. Here we describe the first functioning gene delivery vector based on a human commensal anellovirus.

In this study, we report the development of a self-replicating rescue architecture that enables the trans-complementation of anellovirus-based vector genomes for DNA replication and encapsidation into betatorquevirus capsids. The resulting VLPs bear the hallmark characteristics of anelloviruses. The Anellovectors specifically encapsidate a single-stranded circular genome into a 30-nm non-enveloped particle. This encapsidation is dependent on the ORF1 capsid protein. The ANV.eGFP demonstrates functionality in both *in vitro* and *in vivo* models. Taken all together, these observations suggest that the SATURN platform has been able to reconstitute an anellovirus replication system inside MOLT-4 cells to produce functioning Anellovector.

While directed evolution and machine learning-based design are important tools in the world of viral vector development, the SATURN platform was designed to take advantage of the rich diversity of anelloviruses in the human virome and limit the need to engineer Anellovectors with specific characteristics. As noted by Sandbrink et al, commensal anelloviruses persistently evade the host immune system and hence may not require engineered modifications to overcome anti-vector immunity [Sandbrink 2023]. The development of the SATURN platform has not only enabled the vectorization of the human eye tissue-derived nrVL4619 betatorquevirus, but also the vectorization of two additional members of the *Betatorquevirus* genus. The modular nature of SATURN enables the vectorization of many new anelloviruses that are continuing to be found in tissues throughout the human body. We recently described our rolling circle amplification-based Anelloscope platform that was deployed to discover thousands of anellovirus sequences in an array of human tissue samples [Arze 2021]. The data derived from Anelloscope-based viral sequence hunting provide fertile ground for novel anellovirus sequences that may be vectorizable using the SATURN platform. As we have shown here, nrVL4619-based DNA may be suitable to be used as a universal vector genome that can be replicated and packaged by proteins from a range of unrelated anelloviruses.

The nrVL4619 betatorquevirus was originally sequenced from human ocular tissue [Nawandar 2022]. An Anellovector based on nrVL4619 sequences and produced using the SATURN platform showed penetrance of the Anellovector DNA into the nuclei of cultured, iPSC-derived RPE cells, which resulted in expression of an eGFP payload.

The same Anellovector displayed durable expression over 9 months, as measured by mRNA levels, post-subretinal injection in the murine eye. These findings suggest that the Anellovector retains the ability of the parental virus to transduce its tissue of origin and maintain gene expression over time. The durability of transgene expression is a vital characteristic for any genetic medicine that aims to treat monogenic or polygenic conditions over the lifetime of a patient.

Exploring tropism beyond the eye, we demonstrated the nrVL4619-based Anellovector showed comparable activity to AAV9 following ICV administration in an FVB mouse brain. While additional studies are required to look for markers of inflammation and cell health, the morphology of cells transduced by the Anellovector appeared more typical of healthy neurons when compared to cells transduced by AAV9. Given the reports of AAV9 and other AAV serotype-induced CNS toxicity [Hordeaux 2020, Van Alstyne 2021, Suriano 2021, Guo 2023], the data reported here, though preliminary, suggest that ANV.eGFP may result in less axon degeneration compared to AAV9.eGFP. Anelloviruses have been described as weakly immunogenic and non-pathogenic [Venkataraman 2022, Liou 2022, Kaczorowska 2023, Timmerman 2024]. It will be of great interest to further investigate the comparison between AAV9 and Anellovector-mediated transgene expression in a neuronal system to determine if Anellovector-mediated transgene expression is better tolerated by the host. Altogether, the transduction of cells within the CNS suggests that Anellovectors may have applications in treating CNS-related indications and may offer a favorable profile in terms of vector-derived toxicity.

## METHODS AND MATERIALS

### Cell culture

MOLT-4 cells were obtained from the National Cancer Institute. MOLT-4 cells were maintained at 37°C with 5% CO_2_ in suspension culture in a complete growth medium (Gibco’s RPMI 1640 with 10% fetal bovine serum, supplemented with 1 mM sodium pyruvate, 0.1% Pluronic F-68, and 2 mM L-glutamine) shaking at 100 rpm with >85% relative humidity (RH).

### Small-scale MOLT-4 transfection

MOLT-4 cells were electroporated in 2S buffer (5 mM KCl, 15 mM MgCl_2_, 15 mM HEPES, 150 mM Na_2_HPO_4_ pH 7.2, and 50 mM sodium succinate) using a NEPA electroporator. Cell pellets containing approximately 1E+07 cells were resuspended in 500 µL of 2S buffer along with 50 ug of Cre expression plasmid, 50 ug of ANV.eGFP vector plasmid, and 100 ug of nrVL4619 SRR plasmid, an amount sufficient for 5 cuvettes of the same DNA content. Once electroporation was completed, approximately 300 µl of pre-warmed media was added to each cuvette using a bulb pipette, and the cells were gently mixed to break up clumps. The cell suspension was then transferred to 25 mL of pre-warmed media in a flask. Flasks were incubated for 3 days at 37°C with shaking at 125 rpm before harvest.

### Transfections for *in vivo*-scale vector production

Larger 1L cultures were prepared using a Maxcyte STx electroporator. 3.3 mL of cells at 1.5e8 cells/mL were seeded into a CL1.1 Maxcyte cartridge with 200 µg of ANV.eGFP plasmid, 200 µg of Cre expression plasmid, and 400 µg of nrVL4619 SRR. A Luer-Lock syringe was used to slowly extract the cells from the cartridge, and then cells were slowly dripped directly onto the bottom of the 2L cell culture flask. The cartridge was washed with 3.5 mL of pre-warmed media and then added to the flask. 16.5 mL of pre-warmed media was added to the flask for a 15-minute recovery at 37°C. The flask was then brought up to 1L with pre-warmed media. Then 500mL of the culture was removed and added to an additional 1L flask. Flasks were incubated under standard conditions for 3 days before harvest.

### Viral harvest

MOLT-4 cells were pelleted by centrifugation and then resuspended in lysis buffer containing 50 mM Tris pH 8.0, 0.5% Triton-X100, 100 mM NaCl, 1 × Halt protease inhibitor cocktail (Thermo Fisher Scientific catalog # 78439), 50 mM Tris pH 8.0, and 2 mM MgCl_2_ and 200 U of mSAN nuclease (Arcticzymes catalog #70950-150). Cell lysates were incubated at 37°C for 1 hour. Cell lysates were then clarified at 10,000 × g for 20 minutes at 4°C to pellet cellular debris.

### Iodixanol linear gradients

To generate iodixanol linear gradients, 13 mL of 60% OptiPrep (Sigma-Aldrich catalog # D1556) was layered over with 13 mL of 20% OptiPrep in 26.3-mL polycarbonate tubes.

These tubes were subsequently spun at a 46-degree angle at a speed of 20 rpm for 16 minutes using a Gradient Master (BioComp). The sample tubes were then centrifuged at 347,000 × g and 20°C for 3 hours using a Type 70 Ti rotor (Beckman Coulter). 1-mL fractions were collected from the top of the tube. Each fraction underwent a DNase-protected qPCR assay, following the procedure outlined below.

### DNase-protected qPCR titer assay

5 uL of sample was added to a 20 µL DNase I digestion reaction with 20 U of DNase I. Reactions were incubated at 37°C for 30 minutes in a thermal cycler. 20 µL of Proteinase K digestion mixture (0.1% pluronic F-68, 400 µg Proteinase K, 0.1% SDS, 5 mM EDTA, 10 mM Tris pH8) was added to the reaction and incubated at 55°C for 30 minutes followed by 95°C for 15 minutes in a thermal cycler. The reaction was diluted 1:10 in water for PCR. Samples were then assayed by qPCR on the QuantStudio 5 – Real-Time PCR System (Thermo Fisher, USA) using TaqMan Gene Expression Master Mix (Thermofisher, USA). The sequence detection primers and FAM custom probes that were used in this study were synthesized by Integrated DNA Technologies, USA. Primer and probe sequences are shown in Table 2. All qPCR reactions were performed in triplicate. The standard curve method was used to calculate the amount of viral/vector DNA recovered.

**Table 2:**
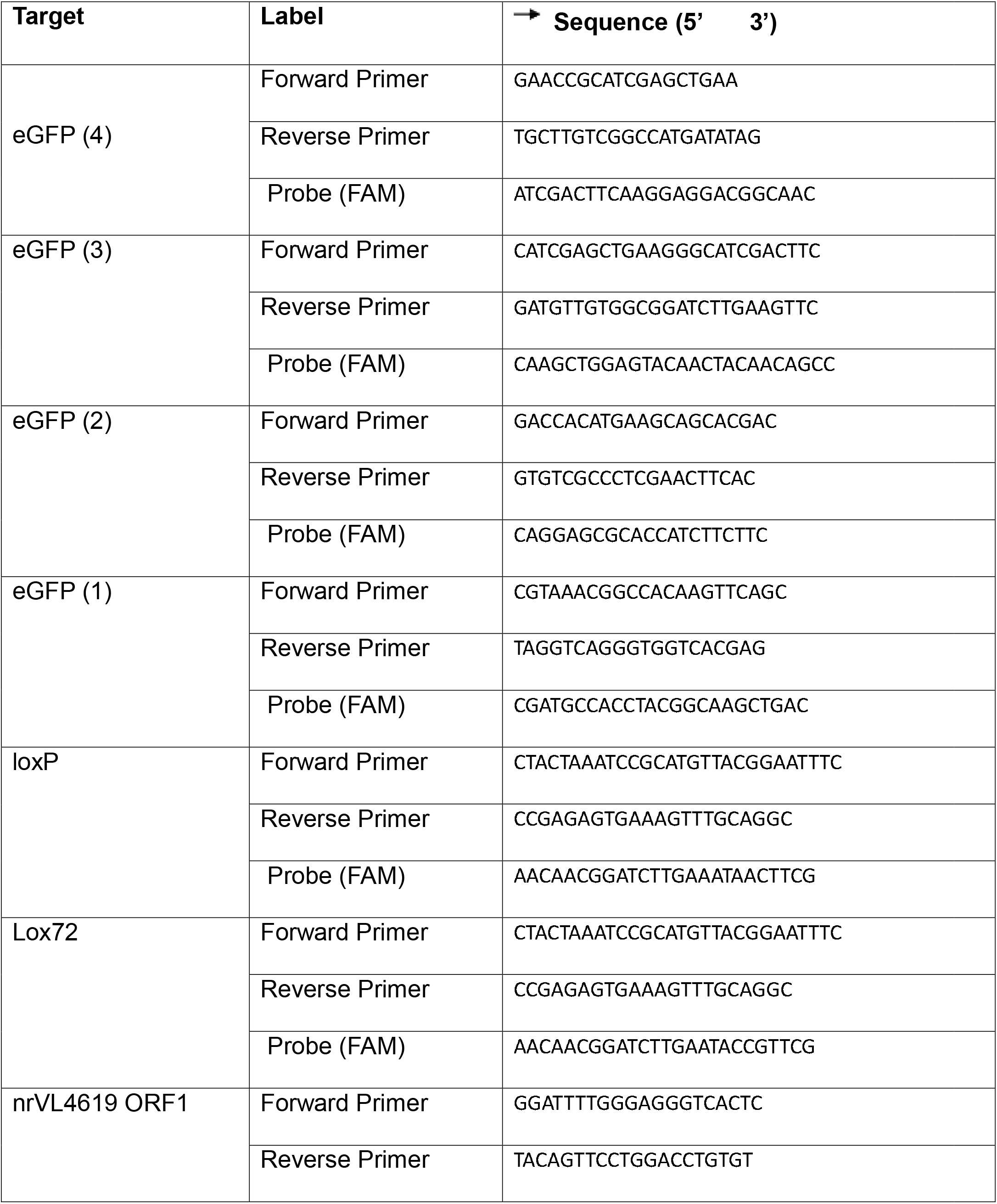
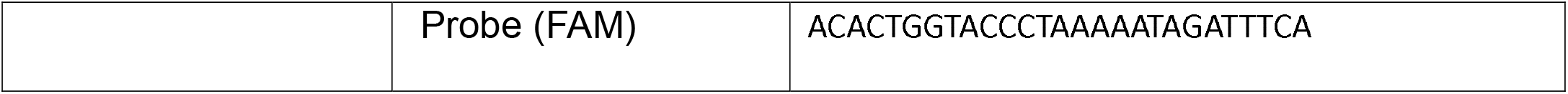
PCR primer and probes.

### Immunoblotting

Cells were harvested 3 days post-electroporation and washed once with PBS followed by lysis in 0.5% Triton-X, 300 mM NaCl, and 50 mM Tris pH 8.0. Genomic DNA was degraded by mSAN, and proteins were denatured using protein sample loading buffer and reducing agent from LI-COR and incubated at 95°C for 10 minutes. Lysates were run on BOLT 12% and 4-12% SDS-PAGE gels for ORF2 and ORF1 immunoblotting, respectively. Transfer to nitrocellulose membranes was performed using the Bio-Rad Trans-Blot Turbo Transfer system. Membranes were blocked for 1 hour in LI-COR Intercept Blocking buffer following by primary antibody incubation (1:1000) overnight at 4°C. GAPDH was detected using an antibody purchased from Cell Signaling (Cat#97166S) and ORF1 and ORF2 were detected using antibodies generated by GenScript. Finally, the membranes were incubated in blocking buffer with IRDye® 800CW and 680CW goat IgG secondary antibodies (1:10,000) from LI-COR for 2 hours and the membranes were imaged using the Bio-Rad ChemiDoc MP imaging system.

### Southern assay

For Southern Blot analysis of replicated viral DNA, the extraction of total nucleic acid from transfected MOLT-4 cells (small scale, as described) was performed using a DNeasy Blood and Tissue Kit (QIAgen catalog # 69504) which includes direct lysis with proteinase K treatment. The DNA was then linearized by digestion at 37oC overnight with a restriction endonuclease unique to the vectorized or wildtype viral DNA of interest respective to each analysis. To digest input plasmid DNA, each sample was also subjected to treatment with DpnI (NEB catalog # R0176) which selectively cleaves methylated GATC sites. After restriction endonuclease treatment, a total of 10µg of DNA per lane was electrophoresed on a 1.0% agarose gel. The electrophoresed DNA was subjected to depurination and denaturation and transferred overnight onto a Hybond-N+ membrane (Cytiva catalog # RPN203B). The membrane was then pre-hybridized for 4h in ULTRAhyb-hybridization buffer (Thermo Fisher Scientific catalog # AM8670) and probed overnight using biotin-labeled oligos unique to the vectorized or wildtype viral DNA of interest. Probes were generated using the BioPrime™ Array CGH Genomic Labeling System (Thermo Fisher Scientific catalog # 18095011). Membranes were blocked for 30m then incubated with IRDye800CW Streptavidin (LI-COR catalong # 926-32230) for 30m and imaged using Bio-Rad ChemiDoc MP Imaging System.

### Electron microscopy

Iodixanol gradient-purified material was adsorbed for ∼1 min to parlodion carbon-coated copper grids, which were rendered hydrophilic by glow discharge at low pressure in air. Grids were washed with three drops of MilliQ water and stained with two drops of 1% uranyl acetate. Electron micrographs were recorded at a nominal magnification of 30,000 × with a JEOL 1400 Flash transmission electron microscope operated at 100 kV and equipped with a NanoSprint43L-MkII CMOS camera system with 43 megapixel sampling region (Advanced Microscopy Techniques).

### Care and use of animals

All mouse procedures were approved and governed by the Ring Therapeutics, Inc Institutional Animal Care and Use Committee (IACUC). All ocular mouse procedures were conducted in compliance with the Association for Research in Vision and Ophthalmology (ARVO) Statement for the Use of Animals in Ophthalmic and Vision Research. Female FVB/NJ (Strain #:001800) and C57Bl/6J (Strain #:00664) mice were purchased from Jackson Laboratories and were acclimated for 72 hours before use in any study. Animals were kept in a 12-hour light (<100Llux)/dark cycle with food and water available ad libitum.

### Subretinal injections□□

Subretinal (SR) injections were performed contralaterally on 9 to 12-week-old C57BI/6J mice. Mice were anesthetized via an intraperitoneal injection of a ketamine/xylazine mixture (Patterson Veterinary Supply, Inc.) at a dose of 100 mg/kg ketamine and 10 mg/kg xylazine. Once anesthetized, pupillary dilation was achieved with one to two drops of a 1% tropicamide/2.5% phenylephrine HCl mixture (Tropi-Phen, Pine Pharmaceuticals). One to two drops of 0.5% proparacaine (McKesson Medical-Surgical, Inc.) were applied to the eye for local anesthetic effect. Subsequently, the animal was positioned in lateral recumbency under a Leica M620 TTS ophthalmic surgical microscope (Leica Microsystems, Inc.). Using a micro scalpel (Surgical Specialties Corporation by Corza Medical), a scleral incision, approximately 0.5 mm long, was made 1 mm posterior to the nasal limbus. A 33-gauge blunt-ended needle (Hamilton Company, Inc.) attached to a 5 μL Hamilton syringe was inserted through the scleral incision, positioned posterior to the lens, and directed towards the temporal retina until resistance was detected. A volume of 1 µL of either vehicle, ANV.eGFP, or ANV.LY2.eGFP vector, containing 0.1% of sodium fluorescein (AK-Fluor 10%, Akorn), was then injected slowly into the subretinal space. The eye was examined under the ophthalmic surgical microscope and the success of the subretinal injection was confirmed by visualizing the fluorescein-containing bleb through the dilated pupil. Only eyes determined to have successful subretinal injections were included in the experiment. Post-procedure, 0.3% tobramycin ophthalmic ointment (Tobrex, McKesson Medical-Surgical Inc.) was applied to each treated eye and the mouse was allowed to recover from the anesthesia prior to being returned to its cage in the housing room.

### SD-OCT system, image acquisition, and analysis

Optical coherence tomography (OCT) retinal images were acquired at 3, 6, and 9 months post-subretinal injection, from independent sets of animals, prior to euthanasia. Animals were anesthetized and eyes were dilated with 1% tropicamide/2.5% phenylephrine HCl (Tropi-Phen, Pine Pharmaceuticals), and corneal protection was provided by 0.3% GenTeal® Gel Eye Drops (Alcon, Inc.). Animals were positioned on a heated staging platform (ALA Scientific Instruments, Inc. and Phoenix-Micron, Inc.) with the Micron IV retinal imaging microscope lens (Phoenix-Micron, Inc.) gently pressed against the eye. Guided by a live bright-field fundus image, the stage and mouse eye were oriented to visualize the subretinal bleb. OCT imaging was performed, and 2D cross-sectional images were captured using InSight v.2 (Voxeleron LLC), an image segmentation software provided by Phoenix Research Labs. The software acquired 20 images and averaged them together to generate a final image for each eye.

Subsequently, the contralateral eye was imaged. Cross-sectional retinal images from each eye were manually measured using InSight v.2 (Voxeleron LLC) image segmentation software provided by Phoenix Research Labs. The internal limiting membrane (ILM) and the outer photoreceptor segment (OS) were manually identified, and the software provided thickness measurements, in microns, across the retinal cross-section image at 1,024 locations. Retinal thickness measurements were averaged, providing a mean retinal thickness for each eye. GraphPad Prism 10.1 was used for statistical analysis. A non-parametric Kolmogorov-Smirnov t-test was used, and groups were considered significantly different if p<0.05.

### Tissue harvest for DNA and RNA extraction

Mouse eyes were harvested at indicated time points following subretinal injections. After enucleation, posterior eye cups (PECs) were separated and collected in 2 mL reinforced tubes (SPEX Sample Prep) containing 5 mM stainless steel beads (Qiagen, LLC).

Samples were flash-frozen on dry ice and stored at −80°C until processing. Brains were cut in half in a sagittal orientation. One half of the brain was processed for DNA and the other half was processed for RNA. Similarly, brains were collected in tubes containing steel beads, flash-frozen and stored at −80°C until ready for homogenization. Frozen tissues were homogenized using Geno/Grinder 2010 (SPEX, SamplePrep, LLC) at 1250 rpm for 30 seconds to 1 minute. Genomic DNA and RNA were isolated from homogenized tissues using the DNEasy Blood and Tissue Kit (Qiagen, LLC) and RNeasy Kit (Qiagen, LLC), respectively, according to the manufacturer’s instructions. DNA concentration was measured using Nanodrop (ThermoFisher, USA) and RNA concentration was measured using Qubit RNA High Sensitivity Assay Kit (ThermoFisher, USA).

### Quantitative PCR analysis

Genomic DNA was assayed by qPCR on the QuantStudio 5 – Real-Time PCR System (Thermo Fisher, USA) using TaqMan Gene Expression Master Mix (Thermofisher, USA). The sequence detection primers and the custom Taqman probe that were used in this study were synthesized by Integrated DNA Technologies, USA. All qPCR reactions were performed in triplicate. The standard curve method was used to calculate the amount of vector DNA, which was normalized with the total amount of genomic DNA for each sample (quantified using Nanodrop as described above). The limit of quantitation (LOQ) was Q=6. GraphPad Prism 10.1 was used for statistical analysis. A non-parametric Mann-Whitney test was used, and groups were considered significantly different if p<0.05.

### One-Step RT-ddPCR analysis

RNA was diluted in nuclease-free water and combined with the reagents from the One Step RT-ddPCR Advanced Kit for Probes (Bio-Rad, USA) and eGFP primer/probe set with final primer concentrations of 900 nM and probe concentrations of 250 nM to measure transgene expression. After the RT-ddPCR reaction setup, each reaction was converted to droplets using the Automated Droplet Generator (Bio-Rad, USA) according to the manufacturer’s instructions. After the droplet generation, the droplets were subjected to endpoint PCR thermocycling with the following cycling conditions: 1 cycle of 48°C for 1 hour for reverse transcription followed by 1 cycle of 95°C for 10 minutes; 40 cycles of 95°C for 30 seconds, 60°C for 1 minute; and 1 cycle of 98°C for 10 minutes and finally a 4°C hold. The cycled plate was then transferred to the QX200 Droplet Reader (Bio-Rad, USA) and analyzed using QX Manager Software (Bio-Rad, USA). GraphPad Prism 10.1 was used for statistical analysis. A non-parametric Mann-Whitney test was used, and groups were considered significantly different if p<0.05.

### AAV vectors

The fluorescent AAV9.eGFP payload was designed in-house at Ring Therapeutics, Inc with ITR flanking elements in order to package the payload into AAV.The AAV9.eGFP plasmid was synthesized by Aldevron (Fargo, North Carolina) and packaged into AAV9 by Packgene Biotech Inc (Houston, Texas) at a titer of 1E13 vgs/ml. Prior to injection, AAV9.eGFP was diluted in sterile 1 × phosphate-buffered saline (PBS) to the titer at 9.1E+10 vg/ml to match with the ANV.eGFP. AAV2-fCMV-eGFP was prepared by Packgene Biotech Inc. and was diluted in sterile 1 × PBS.

### Intracerebroventricular (ICV) injections□□

All animal procedures were in accordance with the Ring Therapeutics, Inc IACUC. FVB/NJ (1800) mice were purchased from Jackson Laboratory and were acclimated for 72 hours before any experiment. Mice were anesthetized with 2%-3% isoflurane and placed with a heating pad on the stereotaxic frame (Neurotar GmbH). Eye ointment was applied to both eyes. After applying aseptic betadine and 70% alcohol, an incision was made to expose the skull. A craniotomy was made by a micro-driller. A 10 µL Hamilton syringe was slowly moved to the target coordinates (lateral ventricles: −0.25 mm in the anteroposterior axis, ±1 mm in the mediolateral axis, and −2.5 mm in the dorsoventral axis). 2 µl of ANV/AAV9.eGFP was injected bilaterally into the left and right lateral ventricles of the brain, making a total injection volume of 4 µl. The injection speed was 0.5 µL/minute. The dose of ANV/AAV9.eGFP was 3.64E+8 vg/mouse. The incision was sutured with absorbable sutures (Suture CT-13.0 coated Vicryl 27”). Each animal received an ICV injection once and a subcutaneous injection of buprenorphine (3.25mg/kg) after the ICV surgery for analgesia on the day of the procedure. Animals were sacrificed 3 weeks after the surgery and brains were collected for tissue processing.

### Immunohistochemistry

Brains were collected and fixed in 4% PFA for 24 hours, then switched to 30% sucrose for an additional 48-72 hours followed by embedding in cryo-preservative media (TissueTek, USA) for sectioning. Cryosectioning was performed on the Leica CM 1850. The sections obtained were 30 µM thick. Slides were air-dried for 20 minutes and then washed with 1 × PBS for 3 x 5min followed by blocking with 10% normal goat serum (Jackson ImmunoReseach Inc.) in TBST for 2 hours at room temperature. Sections were subsequently incubated by primary antibody 1:500 (GFP recombinant rabbit monoclonal antibody; G10362, Life Technologies) for 48 hours at 4°C followed by three PBS washes. The slides were then incubated for 2 hours at room temperature with goat anti-rabbit Alexa 488-conjugated antibody (1:2000; A11008, Life Technologies). Sections were subsequently washed with PBS for 3 x 5 minutes. Slides were mounted under coverslip in Prolong Gold anti-fade reagent with DAPI (Life Technologies.) Stained sections were imaged using EVOS (Invitrogen, Thermo Fisher) and Zeiss LSM900 confocal imaging system, and analyzed using ImageJ software.

### *In vitro* culture and transduction of RPE

iPSC-derived RPE cells were obtained from FujiFilm Cellular Dynamics as a cryopreserved vial and were thawed and cultured according to the manufacturer’s protocol. Briefly, cells were thawed and resuspended in culture media consisting of the MEMα (Thermo Fisher Scientific) supplemented with KnockOut Serum Replacement (Thermo Fisher Scientific), N-2 supplement (Thermo Fisher Scientific), hydrocortisone (Sigma), taurine (Sigma), triiodo-L-thyronine or T3 (Sigma), and gentamicin. RPE cells were spun at 300 g/5 minutes and pellet was resuspended in culture media at 5 × 105 cells/mL. The cells were plated approximately at 1.5 × 10^5^ cell/cm^2^ onto Vitronectin (Thermo Fisher Scientific)-coated glass or plastic surface. RPE cells were maintained at 37°C under 5% CO_2_ in a humidified incubator. The next day a full media change was performed and RPE cells were treated with the Anellovector eGFP (ANV.eGFP) at the desired multiplicity of infection (MOI) (0.9 - 6K). The virus and cell were cultured together for two days, and full media changes were performed on the second day. The RPE cells were maintained in culture and harvested 1,7, and 28 days post-transduction either for immunocytochemistry (ICC) and in-situ hybridization or for running RT-ddPCR. For imaging (ICC and in-situ hybridization), the cells were fixed in 4% paraformaldehyde for 30 minutes and washed three times with DPBS.

### In-situ hybridization

Cells were fixed with 4% paraformaldehyde for 30 minutes at room temperature. Cells were then subject to a dehydration step in 50%, 70%, and 100% ethanol, followed by a rehydration step in 100%, 70%, and 50% ethanol. To rid the cells of free-floating peroxidases, a 10-minute hydrogen peroxide treatment at room temperature was performed. Cells were then immunostained with a GFP (Invitrogen) or Tubulin (Abcam) primary antibody overnight at 4°C. The next day the cells were submerged in 4% paraformaldehyde for 30 minutes at room temperature, followed by a protease III (ACDbio) treatment at a dilution of 1:10 in PBS (Gibco) for 20 minutes. An RNase step was performed using ribonucleases A (25 ug/ml) (Qiagen) and T1 (25 units/ml) (Thermo Scientific) in 1 × Tris-buffered saline containing 0.05% Tween-20 (TBS-T) (Boston Bioproducts) for 30 minutes at 37°C. Following RNase treatment, a dsDNA denaturation step was performed. Cells were subject to 70% formamide (Invitrogen) in 2x saline-sodium citrate (SSC) (Sigma-Aldrich) buffer. Denaturation was performed at 70 °C for 5 minutes. Probe-target hybridization, amplification steps 1-3, and channel development were performed per manufacturer’s protocol from RNAscope Multiplex Fluorescent v2 Assay. Lastly, secondary antibody staining was performed at room temperature for 30 minutes, followed by a 10-minute room temperature DAPI (ACDbio) stain for nuclear visualization. Coverslips were added to slides and left overnight at room temperature to dry. Cells were imaged with a confocal microscope.

### Sequencing

Nucleic acids were extracted from 200 μL viral production samples with a PureLink Viral DNA/RNA kit from Invitrogen (Cat #12280050). Samples were processed according to the manufacturer’s protocol. Samples were eluted in 20 μL of nuclease-free water. Post extraction, DNA was heated for 5 minutes at 95°C before the reaction was quenched on ice. A mastermix containing 2 mmol/L of each dNTP (NEB, Ipswich, MA), 4.5 µg of random hexamers (ThermoFisher, Waltham, MA), and 10 U of DNA polymerase I (NEB) was added to the samples on ice for a final volume of 50 μl. Randomly primed DNA synthesis was performed with a ramp of 0.1°C/second until reaching 37°C, followed by a 1 hour incubation at 37°C. The reaction was stopped with 0.1 mmol/L EDTA. DNA and post-second strand synthesis DNA concentrations were assessed by Qubit. Library preparation of the samples was performed using the Nextera XT (Illumina, San Diego CA, USA) kit, following the manufacturer’s protocol for 1 ng input. Library QC was carried out with D5000 screen tape on an Agilent Tapestation 4200. All libraries were sequenced on the Illumina iSeq 100 platform. Basecalling was performed on Illumina’s BaseSpace cloud computational analysis platform and demultiplexed fastq files were downloaded to local storage. Demultiplexed fastq files were aligned to the payload genome using BWA with default parameters [Li 2013]. Samtools was used to sort, fixmate, and remove duplicates from the resulting BAM files [Danecek 2021]. Alignments were visualized using the Integrative Genomics Viewer (IGV) [Robinson 2011].

## ACKNOWLEDGMENTS

We thank Peter Riebling for support in manuscript writing, and Brian Luque for coordinating submission efforts. Special thanks to Cesar Arze and Jenna Antonucci Johnson for invaluable and meaningful feedback during the preparation of this manuscript.

Sup Figure 1

(A) DNase-protected qPCR assay probing for nrVL4619 WT sequence. Comparison of WT signal in vector samples where iterations of WT nrVL4619 were used as the packaging plasmid. Data presented as mean (n=3). (B) Southern blot probing against eGFP transgene. See Sup Table 1 for lane identities and digestion conditions. The plus and minus signs indicate whether the sample was digested with DpnI to remove input plasmid DNA. Expected band sizes: ANV.eGFP plasmid 5452bp; ANV.eGFP replication intermediate 2839bp. (C) Immunoblots probing for GAPDH and the viral proteins (ORF1 and ORF1/1 top; ORF2/2, and ORF2/3 bottom) from nrVL4619 SRR and ORF1-KO SRR contexts in MOLT-4 cells. Primary antibodies 1:1000, secondary antibodies 1:10,000. The blue arrows denote the expected sizes of the viral proteins. Red arrow denotes the GAPDH loading control. (D) DNase-protected qPCR assay probing for nrVL4619 WT sequences in vector system using nrVL4619 SRR or ORF1-KO SRR to replicate and package ANV.eGFP (n=3). Limit of quantitation represented by dotted line.

Sup Figure 2

(A) Diagram illustrating the Cre recombinase-mediated formation of an ANV.eGFP dsDNA replication intermediate from plasmid DNA and the binding of primers and probes at and around the loxP site of the recombined vector. (B) Electron microscopy of WT nrVL4619 viral particles. 200 nm bar for scale. (C) DNase-protection assay validating the signal from separate egfp, loxP, and lox72 probes.

## REFERENCES

1. Bin Umair M, Akusa FN, Kashif H, Seerat-E-Fatima, Butt F, Azhar M, Munir I, Ahmed M, Khalil W, Sharyar H, Rafique S, Shahid M, Afzal S. Viruses as tools in gene therapy, vaccine development, and cancer treatment. Arch Virol. 2022 Jun;167(6):1387–1404. doi: 10.1007/s00705-022-05432-8. Epub 2022 Apr 24. PMID: 35462594; PMCID: PMC9035288.

2. Epstein AL, Haag-Molkenteller C. Herpes simplex virus gene therapy for dystrophic epidermolysis bullosa (DEB). Cell. 2023 Aug 17;186(17):3523–3523.e1. doi: 10.1016/j.cell.2023.07.031. PMID: 37595560.

3. Charitidis FT, Adabi E, Ho N, Braun AH, Tierney C, Strasser L, Thalheimer FB, Childs L, Bones J, Clarke C, Buchholz CJ. CAR gene delivery by T-cell targeted lentiviral vectors is enhanced by rapamycin induced reduction of antiviral mechanisms. Adv Sci (Weinh). 2023 Dec;10(35):e2302992. doi: 10.1002/advs.202302992. Epub 2023 Oct 30. PMID: 37904721; PMCID: PMC10724389.

4. Zhao Z, Anselmo AC, Mitragotri S. Viral vector-based gene therapies in the clinic. Bioeng Transl Med. 2021 Oct 20;7(1):e10258. doi: 10.1002/btm2.10258. PMID: 35079633; PMCID: PMC8780015.

5. Mendell JR, Al-Zaidy SA, Rodino-Klapac LR, Goodspeed K, Gray SJ, Kay CN, Boye SL, Boye SE, George LA, Salabarria S, Corti M, Byrne BJ, Tremblay JP. Current clinical applications of *in vivo* gene therapy with AAVs. Mol Ther. 2021 Feb 3;29(2):464–488. doi: 10.1016/j.ymthe.2020.12.007. Epub 2020 Dec 10. PMID: 33309881; PMCID: PMC7854298.

6. Kohn DB, Chen YY, Spencer MJ. Successes and challenges in clinical gene therapy. Gene Ther. 2023 Nov;30(10-11):738–746. doi: 10.1038/s41434-023-00390-5. Epub 2023 Nov 8. PMID: 37935854; PMCID: PMC10678346.

7. Wang D, Tai PWL, Gao G. Adeno-associated virus vector as a platform for gene therapy delivery. Nat Rev Drug Discov. 2019 May;18(5):358–378. doi: 10.1038/s41573-019-0012-9. PMID: 30710128; PMCID: PMC6927556.

8. Li X, Le Y, Zhang Z, Nian X, Liu B, Yang X. Viral vector-based gene therapy. Int J Mol Sci. 2023 Apr 23;24(9):7736. doi: 10.3390/ijms24097736. PMID: 37175441; PMCID: PMC10177981.

9. Kuzmin DA, Shutova MV, Johnston NR, Smith OP, Fedorin VV, Kukushkin YS, van der Loo JCM, Johnstone EC. The clinical landscape for AAV gene therapies. Nat Rev Drug Discov. 2021 Mar;20(3):173–174. doi: 10.1038/d41573-021-00017-7. PMID: 33495615.

10. Biagini P. Classification of TTV and related viruses (anelloviruses). Curr Top Microbiol Immunol. 2009;331:21–33. doi: 10.1007/978-3-540-70972-5_2. PMID: 19230555.

11. Butkovic A, Kraberger S, Smeele Z, Martin DP, Schmidlin K, Fontenele RS, Shero MR, Beltran RS, Kirkham AL, Aleamotu’a M, Burns JM, Koonin EV, Varsani A, Krupovic M. Evolution of anelloviruses from a circovirus-like ancestor through gradual augmentation of the jelly-roll capsid protein. Virus Evol. 2023 May 27;9(1):vead035. doi: 10.1093/ve/vead035. PMID: 37325085; PMCID: PMC10266747.

12. Nishizawa T, Okamoto H, Konishi K, Yoshizawa H, Miyakawa Y, Mayumi M. A novel DNA virus (TTV) associated with elevated transaminase levels in posttransfusion hepatitis of unknown etiology. Biochem Biophys Res Commun. 1997 Dec 8;241(1):92–7. doi: 10.1006/bbrc.1997.7765. PMID: 9405239.

13. Fatoba AJ, Adeleke MA. Chicken anemia virus: A deadly pathogen of poultry. Acta Virol. 2019;63(1):19–25. doi: 10.4149/av_2019_110. PMID: 30879309.

14. Virgin HW, Wherry EJ, Ahmed R. Redefining chronic viral infection. Cell. 2009 Jul 10;138(1):30–50. doi: 10.1016/j.cell.2009.06.036. PMID: 19596234.

15. Kaczorowska J, van der Hoek L. Human anelloviruses: diverse, omnipresent and commensal members of the virome. FEMS Microbiol Rev. 2020 May 1;44(3):305–313. doi: 10.1093/femsre/fuaa007. PMID: 32188999; PMCID: PMC7326371.

16. Ninomiya M, Takahashi M, Nishizawa T, Shimosegawa T, Okamoto H. Development of PCR assays with nested primers specific for differential detection of three human anelloviruses and early acquisition of dual or triple infection during infancy. J Clin Microbiol. 2008 Feb;46(2):507–14. doi: 10.1128/JCM.01703-07. Epub 2007 Dec 19. PMID: 18094127; PMCID: PMC2238095.

17. De Vlaminck I, Khush KK, Strehl C, Kohli B, Luikart H, Neff NF, Okamoto J, Snyder TM, Cornfield DN, Nicolls MR, Weill D, Bernstein D, Valantine HA, Quake SR. Temporal response of the human virome to immunosuppression and antiviral therapy. Cell. 2013 Nov 21;155(5):1178–87. doi: 10.1016/j.cell.2013.10.034. PMID: 24267896; PMCID: PMC4098717.

18. Arze CA, Springer S, Dudas G, Patel S, Bhattacharyya A, Swaminathan H, Brugnara C, Delagrave S, Ong T, Kahvejian A, Echelard Y, Weinstein EG, Hajjar RJ, Andersen KG, Yozwiak NL. Global genome analysis reveals a vast and dynamic anellovirus landscape within the human virome. Cell Host Microbe. 2021 Aug 11;29(8):1305–1315.e6. doi: 10.1016/j.chom.2021.07.001. Epub 2021 Jul 27. PMID: 34320399.

19. Varsani A, Opriessnig T, Celer V, Maggi F, Okamoto H, Blomström AL, Cadar D, Harrach B, Biagini P, Kraberger S. Taxonomic update for mammalian anelloviruses (family Anelloviridae). Arch Virol. 2021 Oct;166(10):2943–2953. doi: 10.1007/s00705-021-05192-x. PMID: 34383165.

20. Freer G, Maggi F, Pifferi M, Di Cicco ME, Peroni DG, Pistello M. The virome and its major component, anellovirus, a convoluted system molding human immune defenses and possibly affecting the development of asthma and respiratory diseases in childhood. Front Microbiol. 2018 Apr 10;9:686. doi: 10.3389/fmicb.2018.00686. PMID: 29692764; PMCID: PMC5902699.

21. Liou S, Cohen N, Zhang Y, Acharekar NM, Rodgers H, Islam S, Zeheb L, Pitts J, Arze C, Swaminathan H, Yozwiak N, Ong T, Hajjar RJ, Chang Y, Swanson KA, Delagrave S. Anellovirus structure reveals a mechanism for immune evasion. bioRxiv 2022.07.01.498313.

22. Nawandar DM, Trivedi M, Bounoutas G, Lebo K, Prince C, Scano C, Agarwal N, Ozturk E, Yu J, Arze CA, Bhattacharyya A, Verma D, Thakker P, Cabral J, Liou S-H, Swanson K, Swaminathan H, Diaz F, Mackey A, Chang Y, Ong T, Yozwiak NL, Hajjar RJ, Delagrave S. 2022. Human anelloviruses produced by recombinant expression of synthetic genomes. bioRxiv. 2022.04.28.489885.

23. Hoess R, Abremski K, Irwin S, Kendall M, Mack A. DNA specificity of the Cre recombinase resides in the 25 kDa carboxyl domain of the protein. J Mol Biol. 1990 Dec 20;216(4):873–82. doi: 10.1016/S0022-2836(99)80007-2. PMID: 2266559.

24. Minowada J, Onuma T, Moore GE. Rosette-forming human lymphoid cell lines. I. Establishment and evidence for origin of thymus-derived lymphocytes. J Natl Cancer Inst. 1972 Sep;49(3):891–5. PMID: 4567231.

25. Rao BS, Martin RG. DpnI assay for DNA replication in animal cells: enzyme-resistant material can result from factors not related to replication. Nucleic Acids Res. 1988 May 11;16(9):4171. doi: 10.1093/nar/16.9.4171. PMID: 2836814; PMCID: PMC336596.

26. Wilson VG. Cell culture assay for transient replication of human and animal papillomaviruses. Curr Protoc Microbiol. 2012 Feb;Chapter 14:Unit14B.1. doi: 10.1002/9780471729259.mc14b01s24. PMID: 22307550.

27. Collins KL, Erdile LF, Randall SK, Russo AAR, Simancek PR, Umbricht CB, Virshup DM, Weinberg DH, Wold MS, Kelly TJ. SV40 DNA replication with purified proteins: functional interactions among the initiation proteins. In: Hughes P, Fanning E, Kohiyama M (eds) DNA replication: the regulatory mechanisms. Springer, Berlin, Heidelberg. 10.1007/978-3-642-76988-7_33

28. Albert H, Dale EC, Lee E, Ow DW. Site-specific integration of DNA into wild-type and mutant lox sites placed in the plant genome. Plant J. 1995 Apr;7(4):649–59. doi: 10.1046/j.1365-313x.1995.7040649.x. PMID: 7742860.

29. Galmès J, Li Y, Rajoharison A, Ren L, Dollet S, Richard N, Vernet G, Javouhey E, Wang J, Telles JN, Paranhos-Baccalà G. Potential implication of new torque teno mini viruses in parapneumonic empyema in children. Eur Respir J. 2013 Aug;42(2):470–9. doi: 10.1183/09031936.00107212. Epub 2012 Oct 11. PMID: 23060626; PMCID: PMC3729974.

30. Staurenghi G, Sadda S, Chakravarthy U, Spaide RF; International Nomenclature for Optical Coherence Tomography (IN•OCT) Panel. Proposed lexicon for anatomic landmarks in normal posterior segment spectral-domain optical coherence tomography: the IN•OCT consensus. Ophthalmology. 2014 Aug;121(8):1572–8. doi: 10.1016/j.ophtha.2014.02.023. Epub 2014 Apr 19. PMID: 24755005.

31. Zeppieri M, Marsili S, Enaholo ES, Shuaibu AO, Uwagboe N, Salati C, Spadea L, Musa M. Optical Coherence Tomography (OCT): A Brief Look at the Uses and Technological Evolution of Ophthalmology. Medicina (Kaunas). 2023 Dec 3;59(12):2114. Doi: 10.3390/medicina59122114. PMID: 38138217; PMCID: PMC10744394.

32. Wang C, Pan C, Yong H, Wang F, Bo T, Zhao Y, Ma B, He W, Li M. Emerging non-viral vectors for gene delivery. J Nanobiotechnology. 2023 Aug 17;21(1):272. doi: 10.1186/s12951-023-02044-5. PMID: 37592351; PMCID: PMC10433663.

33. Rajawat YS, Humbert O, Kiem HP. In-Vivo Gene Therapy with Foamy Virus Vectors. Viruses. 2019 Nov 23;11(12):1091. doi: 10.3390/v11121091. PMID: 31771194; PMCID: PMC6950547.

34. Arabi F, Mansouri V, Ahmadbeigi N. Gene therapy clinical trials, where do we go? An overview. Biomed Pharmacother. 2022 Sep;153:113324. doi: 10.1016/j.biopha.2022.113324. Epub 2022 Jun 29. PMID: 35779421.

35. Shen W, Liu S, Ou L. rAAV immunogenicity, toxicity, and durability in 255 clinical trials: A meta-analysis. Front Immunol. 2022 Oct 27;13:1001263. doi: 10.3389/fimmu.2022.1001263. Erratum in: Front Immunol. 2023 Jan 18;13:1104646. PMID: 36389770; PMCID: PMC9647052.

36. Srivastava A. Rationale and strategies for the development of safe and effective optimized AAV vectors for human gene therapy. Mol Ther Nucleic Acids. 2023 May 17;32:949–959. doi: 10.1016/j.omtn.2023.05.014. PMID: 37293185; PMCID: PMC10244667.

37. Sandbrink JB, Alley EC, Watson MC, Koblentz GD, Esvelt KM. Insidious Insights: Implications of viral vector engineering for pathogen enhancement. Gene Ther. 2023 May;30(5):407–410. doi: 10.1038/s41434-021-00312-3. Epub 2022 Mar 10. PMID: 35264741; PMCID: PMC10191845.

38. Hordeaux J, Buza EL, Dyer C, Goode T, Mitchell TW, Richman L, Denton N, Hinderer C, Katz N, Schmid R, Miller R, Choudhury GR, Horiuchi M, Nambiar K, Yan H, Li M, Wilson JM. Adeno-Associated Virus-Induced Dorsal Root Ganglion Pathology. Hum Gene Ther. 2020 Aug;31(15-16):808–818. Doi: 10.1089/hum.2020.167. Epub 2020 Jul 31. Erratum in: Hum Gene Ther. 2021 Jul 20;: PMID: 32845779.

39. Van Alstyne M, Tattoli I, Delestrée N, Recinos Y, Workman E, Shihabuddin LS, Zhang C, Mentis GZ, Pellizzoni L. Gain of toxic function by long-term AAV9-mediated SMN overexpression in the sensorimotor circuit. Nat Neurosci. 2021 Jul;24(7):930–940. doi: 10.1038/s41593-021-00827-3. Epub 2021 Apr 1. PMID: 33795885; PMCID: PMC8254787.

40. Suriano CM, Verpeut JL, Kumar N, Ma J, Jung C, Boulanger LM. Adeno-associated virus (AAV) reduces cortical dendritic complexity in a TLR9-dependent manner. bioRxiv 2021.09.28.462148.

41. Guo Y, Chen J, Ji W, Xu L, Xie Y, He S, Lai C, Hou K, Li Z, Chen G, Wu Z. High-titer AAV disrupts cerebrovascular integrity and induces lymphocyte infiltration in adult mouse brain. Mol Ther Methods Clin Dev. 2023 Aug 28;31:101102. Doi: 10.1016/j.omtm.2023.08.021. PMID: 37753218; PMCID: PMC10518493.

42. Venkataraman T, Swaminathan H, Arze CA, Jacobo SM, Bhattacharyya A, David T, Nawandar DM, Delagrave S, Mani V, Yozwiak NL, Larman HB. Comprehensive profiling of antibody responses to the human anellome using programmable phage display. Cell Rep. 2022 Dec 20;41(12):111754. doi: 10.1016/j.celrep.2022.111754. PMID: 36543141.

43. Kaczorowska J, Timmerman AL, Deijs M, Kinsella CM, Bakker M, van der Hoek L. Anellovirus evolution during long-term chronic infection. Virus Evol. 2023 Jan 5;9(1):vead001. Doi: 10.1093/ve/vead001. PMID: 36726484; PMCID: PMC9885978.

44. Timmerman AL, Schönert ALM, van der Hoek L. Anelloviruses versus human immunity: how do we control these viruses? FEMS Microbiol Rev. 2024 Jan 12;48(1):fuae005. doi: 10.1093/femsre/fuae005. PMID: 38337179; PMCID: PMC10883694.

45. Li H. Aligning sequence reads, clone sequences and assembly contigs with BWA-MEM. arXiv. 2013; doi: 10.48550/arxiv.1303.3997

46. Danecek P, Bonfield JK, Liddle J, Marshall J, Ohan V, Pollard MO, Whitwham A, Keane T, McCarthy SA, Davies RM, Li H. Twelve years of SAMtools and BCFtools. GigaScience. 2021;10(2):giab008. PMCID: PMC7931819

47. Robinson JT, Thorvaldsdóttir H, Winckler W, Guttman M, Lander ES, Getz G, Mesirov JP. Integrative genomics viewer. Nat Biotechnol. 2011;29(1):24–26. PMCID: PMC3346182

