## Supplementary figures and images for "A novel functional gene delivery platform based on a commensal human anellovirus demonstrates transduction in multiple tissue types"

### Supplemental Figure 1

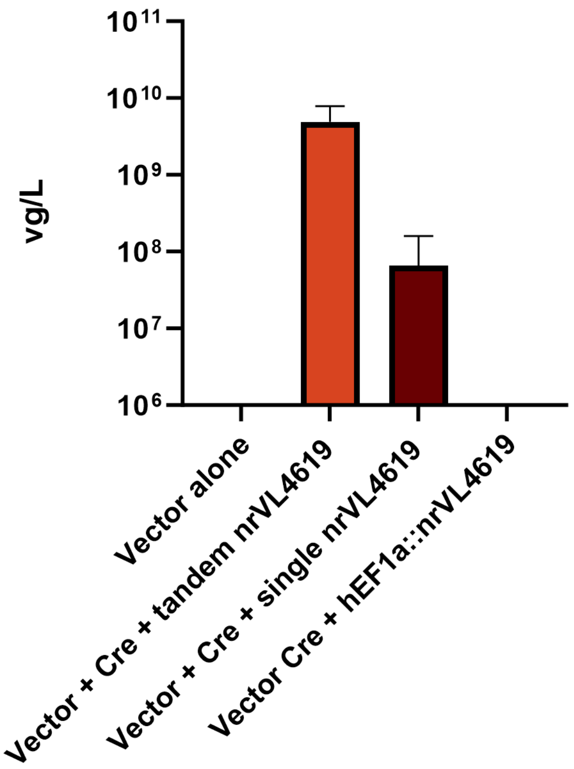

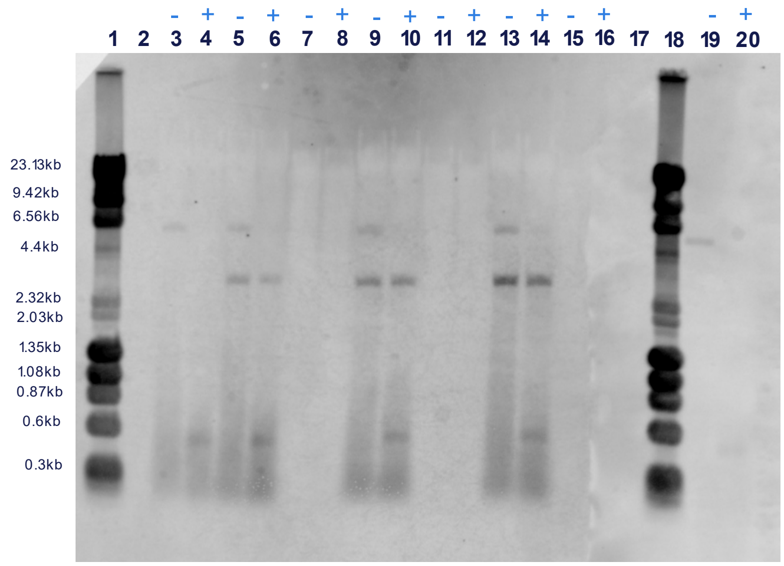

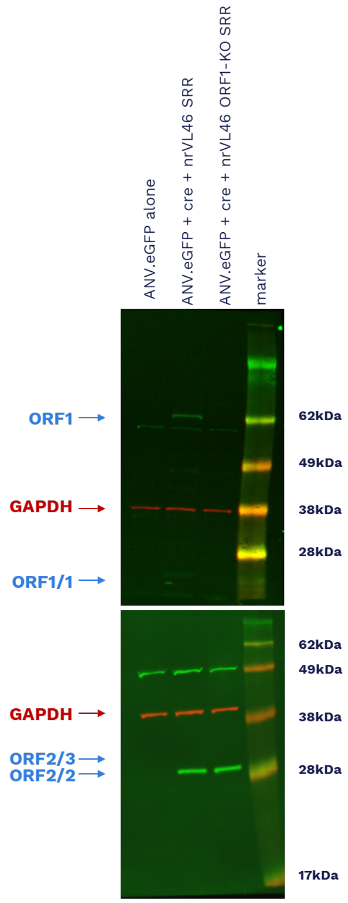

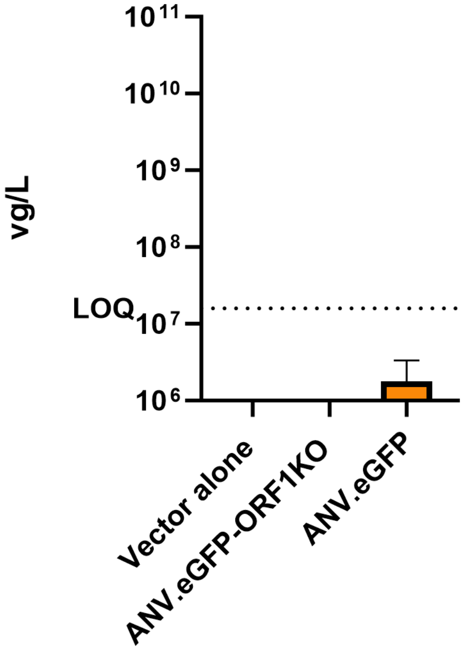

### Supplemental Figure 2

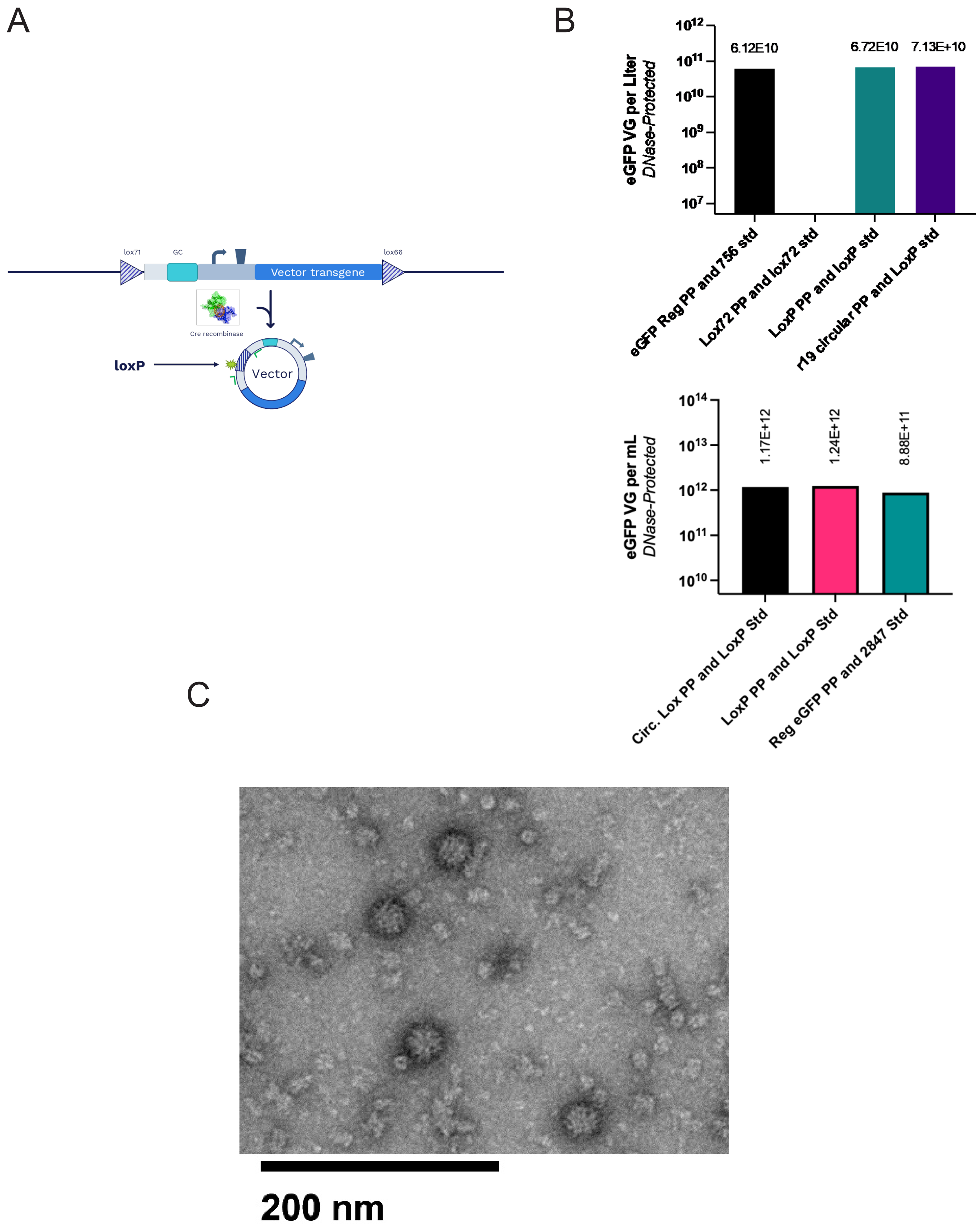
