## Supplemental Table 1 for "A novel functional gene delivery platform based on a commensal human anellovirus demonstrates transduction in multiple tissue types"

### Probe: eGFP

| Lane | Condition | Enzyme(s) |
|---|---|---|
| **1** | Mixed Ladder: λ DNA-HindIII Digest + ΦX174 DNA-HaeIII Digest | |
| **2** | empty | |
| **3** | ANV.eGFP alone | AgeI-HF |
| **4** | ANV.eGFP alone | AgeI-HF **+ DpnI** |
| **5** | ANV.eGFP + Cre + nrVL4619 Tandem | AgeI-HF |
| **6** | ANV.eGFP + Cre + nrVL4619 Tandem | AgeI-HF **+ DpnI** |
| **7** | nrVL4619 Tandem alone | EcoRV-HF |
| **8** | nrVL4619 Tandem alone | EcoRV-HF **+ DpnI** |
| **9** | ANV.eGFP + Cre + nrVL4619 single copy | AgeI-HF |
| **10** | ANV.eGFP + Cre + nrVL4619 single copy | AgeI-HF **+ DpnI** |
| **11** | nrVL4619 single copy alone | EcoRV-HF |
| **12** | nrVL4619 single copy alone | EcoRV-HF **+ DpnI** |
| **13** | ANV.eGFP + Cre + hEF1a-nrVL4619 CDS | AgeI-HF |
| **14** | ANV.eGFP + Cre + hEF1a-nrVL4619 CDS | AgeI-HF **+ DpnI** |
| **15** | hEF1a-nrVL4619 CDS alone | EcoRV-HF |
| **16** | hEF1a-nrVL4619 CDS alone | EcoRV-HF **+ DpnI** |
| **17** | empty | |
| **18** | Mixed Ladder: λ DNA-HindIII Digest + ΦX174 DNA-HaeIII Digest | |
| **19** | 1e+09 copies of ANV.eGFP plasmid (control plasmid digest) | AgeI-HF |
| **20** | 1e+09 copies of ANV.eGFP plasmid (control plasmid digest) | AgeI-HF + DpnI |
